# Explainable Machine Learning Reveals the Role of the Breast Tumor Microenvironment in Neoadjuvant Chemotherapy Outcome

**DOI:** 10.1101/2023.09.07.556655

**Authors:** Youness Azimzade, Mads Haugland Haugen, Xavier Tekpli, Chloé B. Steen, Thomas Fleischer, David Kilburn, Hongli Ma, Eivind Valen Egeland, Gordon Mills, Olav Engebraaten, Vessela N. Kristensen, Arnoldo Frigessi, Alvaro Köhn-Luque

## Abstract

Recent advancements in single-cell RNA sequencing (scRNA-seq) have enabled the identification of phenotypic diversity within breast tumor tissues. However, the contribution of these cell phenotypes to tumor biology and treatment response has remained less understood. This is primarily due to the limited number of available samples and the inherent heterogeneity of breast tumors. To address this limitation, we leverage a state-of-the-art scRNA-seq atlas and employ CIBER-SORTx to estimate cell phenotype fractions by de-convolving bulk expression profiles in more than 2000 samples from patients who have undergone Neoadjuvant Chemotherapy (NAC). We introduce a pipeline based on explainable Machine Learning (XML) to robustly explore the associations between different cell phenotype fractions and the response to NAC in the general population as well as different subtypes of breast tumors. By comparing tumor subtypes, we observe that multiple cell types exhibit a distinct association with pCR within each subtype. Specifically, Dendritic cells (DCs) exhibit a negative association with pathological Complete Response (pCR) in Estrogen Receptor positive, ER+, (Luminal A/B) tumors, while showing a positive association with pCR in ER-(Basal-like/HER2-enriched) tumors. Analysis of new spatial cyclic immunofluorescence data and publicly available imaging mass cytometry data showed significant differences in the spatial distribution of DCs between ER subtypes. These variations underscore disparities in the engagement of DCs within the tumor microenvironment (TME), potentially driving their divergent associations with pCR across tumor subtypes. Overall, our findings on 28 different cell types provide a comprehensive understanding of the role played by cellular components of the TME in NAC outcomes. They also highlight directions for further experimental investigations at a mechanistic level.

## 1 Introduction

NAC is the established treatment approach for patients with large breast tumors, involving the administration of chemotherapy drugs prior to surgical removal of the tumor [1]. Following the completion of NAC, pCR which refers to the complete elimination of cancer cells in the breast and auxiliary lymph nodes, serves as a highly favorable prognostic biomarker [2, 3]. Consequently, gaining a better understanding of pCR in response to NAC is not only of biological interest but also holds significant clinical relevance [4, 5].

Breast tumors are complex ecosystems comprising cancer cells, normal epithelial cells, immune cells and stromal cells [6, 7]. Recent advancements in scRNA-seq of tumor tissue samples have provided detailed insights into gene expression patterns, revealing the heterogeneity within tumor cells as well as within the surrounding cellular milieu, collectively referred to as the TME [8]. Despite the progress, the contribution of such cell phenotypes to tumor biology and response to NAC remains poorly understood and still limited to well-known aspects such as the role of tumor proliferation and immune infiltration [4].

The availability of high-resolution scRNA-seq data needed for comprehensive analysis is currently limited to a small number of samples [8–12]. Consequently, the statistical power required for a thorough analysis of the diverse range of cell types present in tissue samples is still lacking. Moreover, breast tumors exhibit substantial heterogeneity and encompass various subtypes characterized by the status of Estrogen Receptor (ER), Progesterone Receptor (PR) and Human Epidermal Growth Factor Receptor 2 (HER2) [13], or by their expression profile using the PAM50 method [14]. Each of these subtypes displays a distinct TME [15] and exhibits significantly different response rates to chemotherapy [16]. The combination of these factors limits our current understanding and makes significant progress in the near future challenging, emphasizing the need for alternative approaches.

One such approach is deciphering cell phenotypes within bulk gene expression profile (bGEP) through deconvolution based on marker genes [17, 18] for which a wide range of methods have been developed [19]. The large number of samples with available bGEP can potentially provide the necessary statistical power for a proper analysis [20, 21]. However, traditional statistical models often encounter difficulties when dealing with complex, highly non-linear problems with intricate interactions, such as the role of cell types in pCR. Conversely, machine learning (ML) models excel in capturing such complex patterns and have, as a result, revolutionized cancer research by introducing transformative opportunities in diagnostic, prognostic and foundational studies [22–24]. Advances in XML [25, 26] have further improved the utility of ML methods by identifying and quantifying the role of the complex patterns that contribute to individual predictions [27].

In this paper, we first establish a Signature Matrix (SM), a cell-specific expression profile for all cell types expected to be present in a tumor tissue sample, utilizing a scRNA-seq atlas of human breast cancers [8]. We leverage this SM to estimate cell fractions from bGEPs of more than 2000 tumor samples using CIBERSORTx [28]. The samples were then divided into two cohorts, namely discovery and the validation cohorts. We then present a novel pipeline based on XML which enables filtering out prediction-specific patterns. Within this pipeline, we introduce “pCR Score” as a robust metric for the association of fraction of cell types with the probability of getting pCR. The combined utilization of a large number of samples and the high resolution of the tumor tissue, along with XML, allows us to effectively calculate pCR Score in the general population (GP), as well as in different breast cancer subtypes. Inspired by our discoveries, we delve deeper into the analysis of scRNA-seq and spatial omics data of DCs in order to uncover the underlying factors contributing to their contrasting association with pCR across tumor subtypes.

## 2 Results

### 2.1 pCR Scores in the general population

To deconvolve bulk gene expression profiles, we utilized a scRNA-seq breast cancer atlas comprising 100,000 cells collected from 26 patient samples [8]. We constructed an SM by selecting 28 different cell types, including subsets of B cells, T cells, cancer-associated fibroblasts (CAFs), perivascular-like cells (PVLs), plasmablasts, myeloids, endothelial cells, normal epithelial cells and cancer cells. Cancer cells were divided into seven recurrent Gene Modules (GenMod1-7), based on similarities in their expression profiles. Each GenMod was defined with 200 genes and exhibited a high correlation with one or a few well-known biological pathways as follows: GenMod1: Jun.Fos, ER, RTKs, p53; GenMod2: Myc, OxPhos; GenMod3: EMT, IFN, Complement; GenMod4: Proliferation; GenMod5: Classical ER; GenMod6: Diverse; GenMod7: RTKs [8].

Our study included breast tumor samples that met the following inclusion criteria: (1) they underwent NAC, (2) gene expression microarray data was available at baseline and (3) the response to treatment was assessed as either pCR or residual disease (RD). We identified 15 datasets with a total of 2,007 samples (485 pCR, 1,522 RD). Patients in these datasets were treated mostly with Taxane (14 datasets, 1910 patients) alongside arrays of other drugs such as Fluorouracil (9 datasets, 1156 patients), Cyclophosphamide (8 datasets, 1059 patients), Capecitabine (5 datasets, 816 patients), etc. Apart from gene expression profiles, we were able to retrieve two features in these datasets: ER status and PAM50 subtypes. These features were included as covariates in our ML models.

Each dataset was log2 normalized if needed and then cell type fractions were calculated using CIBERSORTx. Subsequently, we randomly divided the datasets into two sets with a similar number of samples: the discovery (9 datasets including 1,009 samples) and validation (6 datasets including 998 samples) cohorts (see Methods and SI for details). Then, we applied steps II and III of our pipeline to calculate pCR Scores (refer to Fig. 1 and Methods for a precise description of the analysis pipeline). Fig. 2 illustrates the final result of our pipeline applied to the general population. Dividing the data into two parts and developing separate ML models for each part, followed by validation (steps II and III), increased the generalizability of the proposed pCR Scores.

**Fig. 1:**
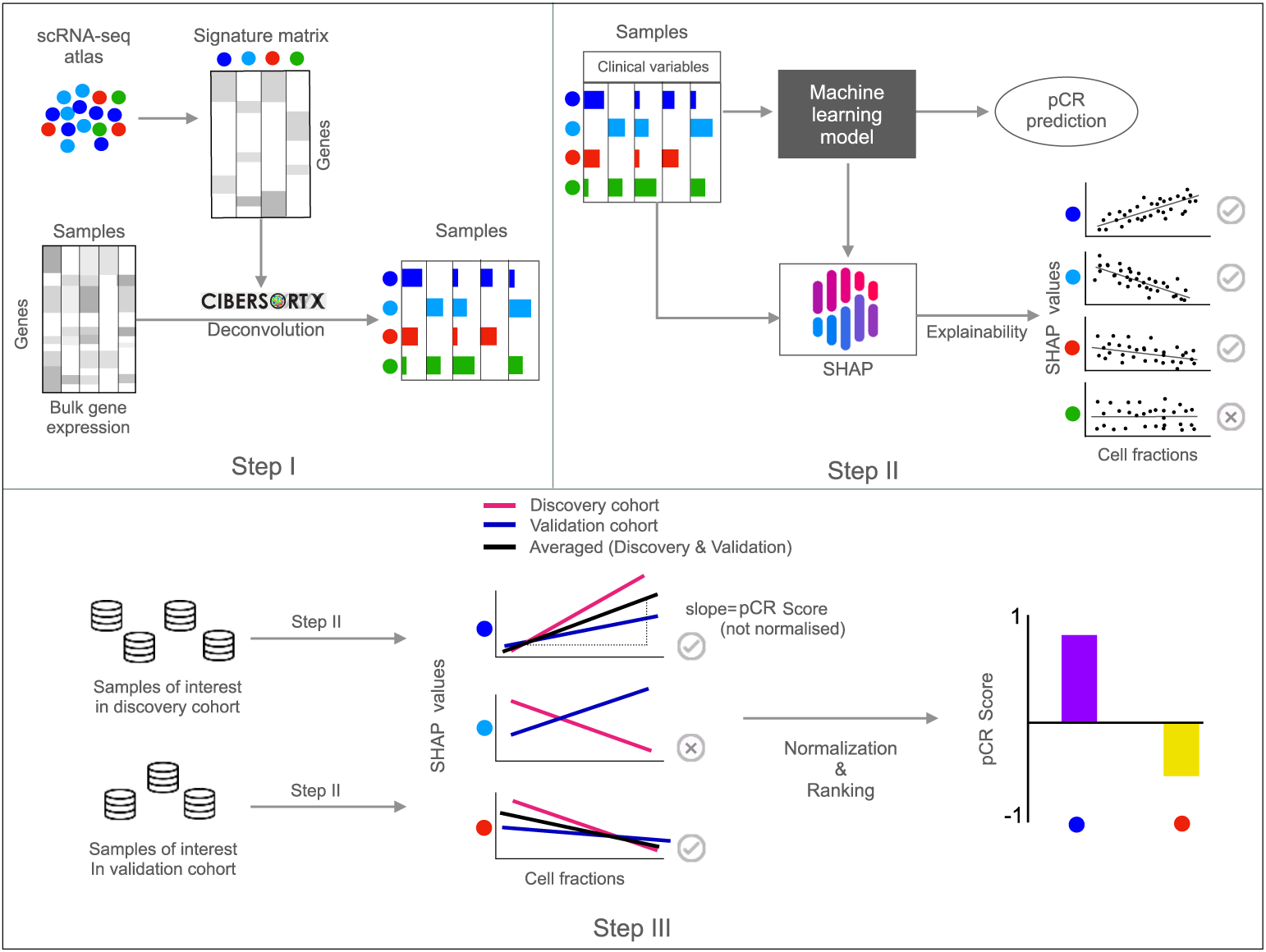
a) We define a high-resolution Signature Matrix (SM) for breast tumor tissue and validate its performance on pseudobulk data (see Supplementary Information (SI)). Cell types in colors. This SM is then used to deconvolve bulk gene expression data (Step I). b) For a given group of samples from the discovery or the validation cohort, we utilize cell fractions, ER status and PAM50 subtype to train an ML model that predicts pCR. We subsequently calculate SHAP values for all cell types in the given samples. Cell types that do not exhibit a clear association with pCR are excluded. c) In step III, we compare the SHAP values vs. cell fractions in the discovery and the validation cohorts. Cell types with consistent associations in both cohorts (either both negative or both positive) are considered further. For these cell types, we calculate the average slope of the SHAP value vs. cell fraction, normalize it and obtain the pCR Score of each cell type. Finally, cell types are ranked based on their pCR Scores.

**Fig. 2:**
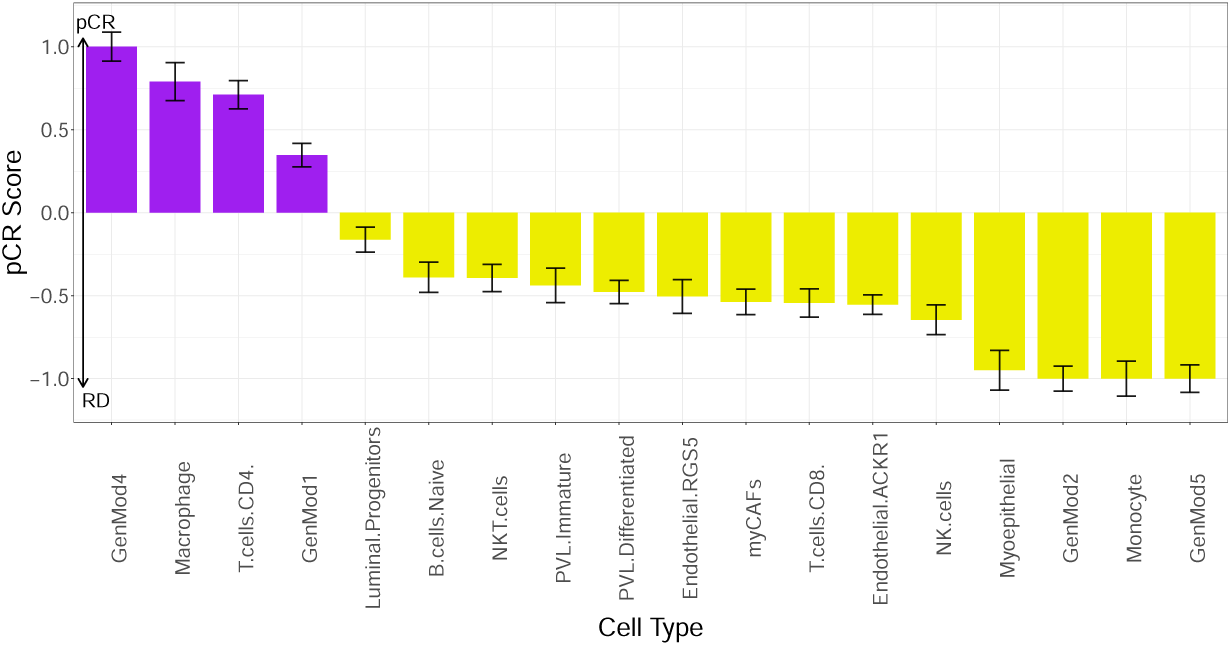
Ranked pCR Scores of cell types calculated in the general population (here and throughout this paper, error bars represent a 0.999 confidence interval). A combination of cancer cells (GenMod4, GenMod1, GenMod2 and GenMod5), immune cells (CD4 T cells, macrophages, naive B cells, NKT cells, CD8 T cells and NK cells), stromal cells (immature PVLs, differentiated PVLs, RGS5 endothelials, ACKR1 endothelials and myCAFs) and normal epithelial cells (luminal progenitors, myoepithelials) exhibit non-zero pCR Scores.

#### Cell types with positive pCR Scores in the general population

Among the cancer cell types, GenMod4 and GenMod1 exhibit positive pCR Scores. The association of GenMod4 aligns with simple t-test results, where samples with pCR displayed significantly higher fractions of GenMod4 compared to samples with RD (see Suppl. Fig. S4). Immune cells including macrophages and CD4 T cells also demonstrate positive pCR Scores.

#### Cell types with negative pCR Scores in the general population

Cancer cell types GenMod5 and GenMod2 exhibit negative pCR Scores. Consistently, these cell types are found in significantly lower proportions within samples that achieve pCR compared to those with RD (see Suppl. Fig. S4). Among immune cells, the following cell types are negatively associated with pCR: monocytes, Natural Killer (NK) cells, CD8 T cells, Natural Killer T (NKT) cells and naive B cells. Within stromal cells, the following cell types exhibit a negative association with pCR: ACKR1 endothelials, cancer-associated fibroblasts with myofibroblast characteristics (myCAFs), RGS5 endothelials, differentiated PVLs and immature PVLs. From normal epithelial cells, myoepithelials and luminal progenitors show negative pCR Scores.

While we identified non-zero pCR Scores for many cell types, it is important to note that tumors consist of different subtypes that respond differently to treatment [13]. For instance, ER+ samples contain significantly higher fractions of GenMod5 (refer to Suppl. Fig. S5). Furthermore, in our data, the pCR rate is 13 % for ER+ samples and 37 % for ER-samples. Therefore, when considering the general population, the observed negative pCR Score for GenMod5, despite the ER status (and PAM50 sub-type) being included as covariates in the models, could be influenced by a combination of these two factors. Consequently, it is necessary to separately analyze ER- and ER+ samples to determine whether the observed negative pCR Score solely reflects the differences between ER subtypes or if GenMod5 also acts as an independent feature that regulates treatment response. This is addressed in the following sections.

### 2.2 pCR Scores in ER status subtypes

Among the samples gathered here, 55 % are ER+ and the remaining 45 % are ER-. In addition to the difference in pCR rates, various cell types have significantly different fractions between ER- and ER+ subtypes (refer to Suppl. Fig. S5). However, within each ER subtype, no cell type shows a significant difference in samples with pCR vs. RD (t-test, see Suppl. Fig. S6). This further underscores the opportunity to utilize ML methods to unravel the role of cell types in treatment response.

To calculate pCR Scores in each ER status subtype, we selected ER+ (and ER-) samples from the discovery and validation cohorts and followed steps II and III of the pipeline as described earlier (see Fig. 1). The normalized and sorted pCR Scores for ER+ and ER- samples are displayed in Figs 3 (a) and (b), respectively. The cell types are also ordered as in Fig. 2 (c) based on their pCR Scores in the general population, enabling an easier comparison between ER subtypes as well as with the general population (see Fig. 3 (c)).

**Fig. 3:**
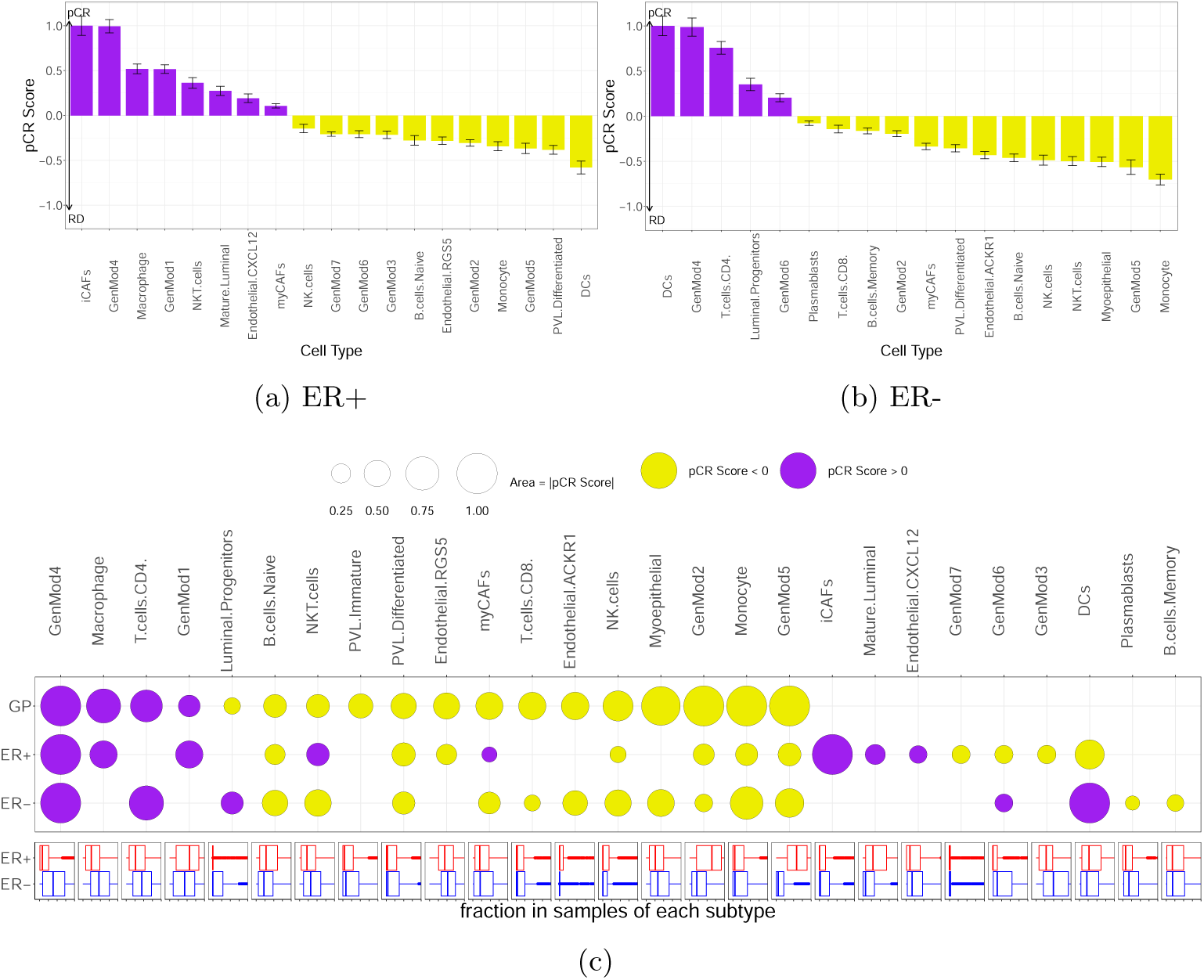
pCR Scores were calculated separately for ER+ samples (a) and ER- (b) samples. c) The pCR Scores were ranked based on the general population (GP). The lower panel displays the normalized fraction of corresponding cell types in ER+ and ER-samples (refer to the SI for details). GenMod4, naive B cells, differentiated PVLs, NK cells, GenMod2, monocytes and GenMod5 exhibit pCR Scores with the same polarities in both subtypes. However, some cell types show high pCR Scores only in one subtype. More interestingly, the polarity of pCR Scores (favoring or disfavouring pCR) for DCs, myCAFs, NKT cells and GenMod6 change across ER subtypes. Additionally, pCR Scores do not seem to depend on the fractions present in subtypes.

#### Cell types with the same polarity of pCR Scores across ER subtypes

Among cancer cell types, GenMod4, GenMod2 and GenMod5 exhibit pCR Scores with the same polarity (sign) in both ER subtypes, despite significant differences in their fractions in ER-vs. ER+ (see Suppl. Fig. S5). Additionally, naive B cells, differentiated PVLs and monocytes demonstrate pCR Scores with the same polarity in both ER subtypes. This consistency suggests that while there are many differences between ER+ and ER-tumors, certain treatment response processes remain the same in both subtypes. It is worth mentioning that the non-zero pCR Score of GenMod5 in both ER subtypes indicates its independent association with pCR even within ER subtypes.

#### Cell types showing pCR Scores only in one of the ER subtypes

Despite the similarities, several cell types exert their effects in an ER status-dependent manner. Macrophages, GenMod1, immature PVLs, RGS5 endothelials, inflammatory CAFs (iCAFS), mature luminals, CXCL12 endothelials, GenMod7 and GenMod3 show non-zero pCR Scores only in the ER+ subtype. On the other hand, CD4 T cells, luminal progenitors, NKT cells, CD8 T cells, ACKR1 endothelials, NK cells, myoepithelials, plasmablasts and memory B cells show non-zero pCR Scores only in ER-samples.

Despite the larger statistical power, not all of these associations were identified in the analysis of the general population. For example, the association of luminal progenitors and immature PVLs with pCR is opposite to what was identified for the general population. This further demonstrates how the associations of cell types with treatment response in specific ER subtypes can differ from the general population, indicating the presence of interactions.

#### Cell types exhibiting opposite polarities in pCR Scores across ER sub-types

Furthermore, for DCs, myCAFs, NKT and GenMod6 the polarity of the pCR Scores change depending on ER subtype. In particular, while the fraction of DCs is not associated with pCR in the general population, in ER+, they are negatively associated with pCR, while in ER-, they are positively associated with pCR. NKT cells and myCAFs show positive pCR Scores in ER+ samples and negative pCR Scores in ER-samples.

### 2.3 pCR Score in PAM50 subtypes

PAM50 subtypes include four major groups as Luminal A, Luminal B, Basal-like and HER2-enriched which represent 29.3 %, 23.1 %, 27.5 % and 12.5 % of our samples, respectively (Normal-like subtype with 7.5 % of samples are not explored here). Each PAM50 subtype exhibits a distinct pCR rate (in our samples Luminal A, Luminal B, Basal-like and HER2-enriched have pCR rates of 7.6 %, 20.4 %, 37.4 % and 39.3 %, respectively). Their microenvironment has a unique combination of cells (see Suppl. Fig. S7), however, within each subtype, no cell type shows a significant difference between the two response groups.

To analyze the PAM50 subtypes, we followed steps II and III of our pipeline separately for each subtype. We calculated distinct pCR Scores for Luminal A (Fig. 4 (a)), Luminal B (Fig. 4 (b)), Basal-like (Fig. 4 (c)) and HER2-enriched (Fig. 4 (d)) sub-types. Additionally, Cell types are ranked based on their pCR Scores in the general population in Fig. 4 (e). Each PAM50 subtype is associated with a unique set of cell types in response, indicating not only a different microenvironment but also different roles for each cell type in NAC response.

**Fig. 4:**
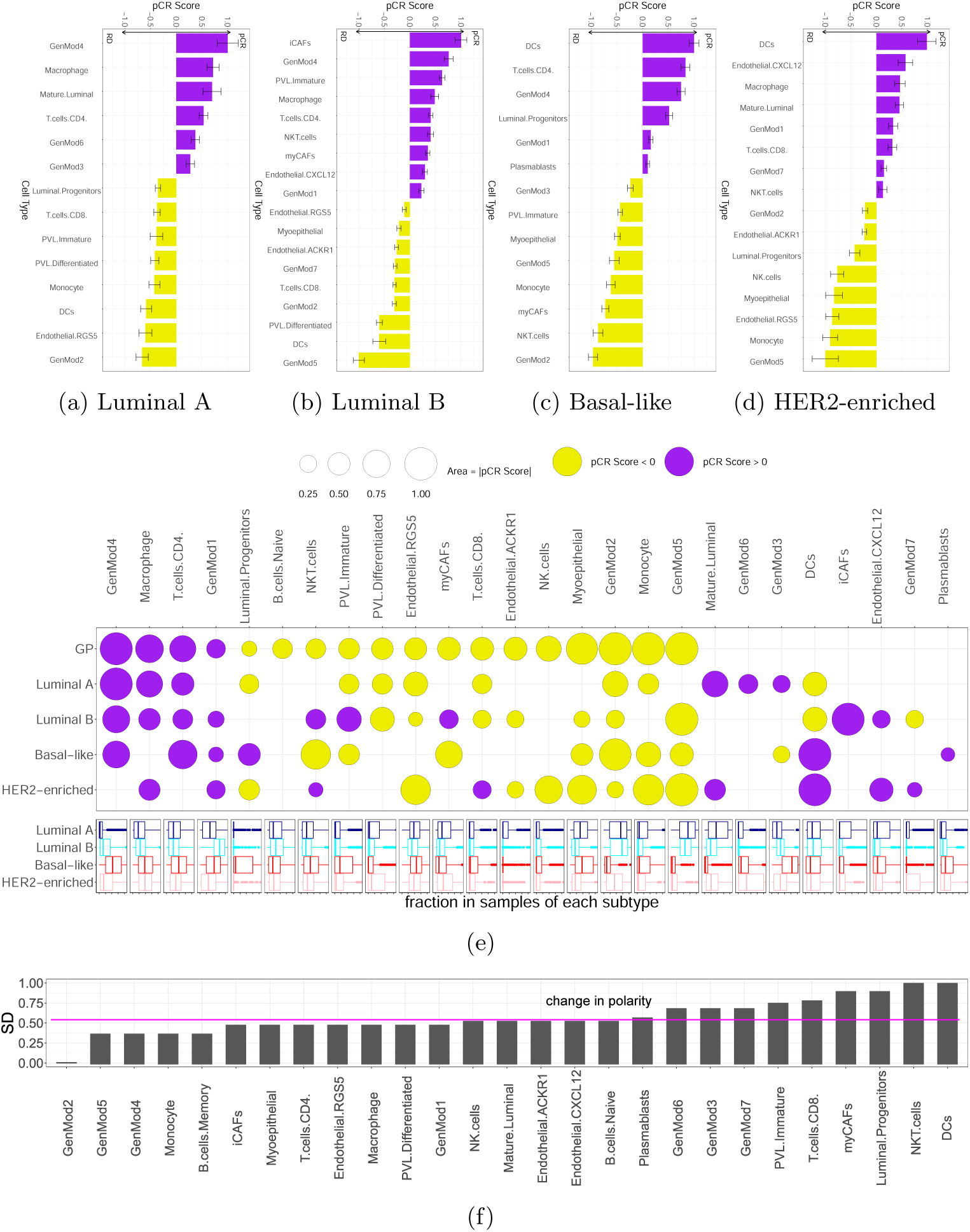
Ranked pCR Scores for Luminal A (a), Luminal B (b), Basal-like (c) and HER2-enriched (d) subtypes. e) pCR Scores ranked based on the general population. The lower panel displays the normalized fraction of the corresponding cell types for PAM50 subtypes. While many cell types share the same polarity of pCR Score across PAM50 subtypes, DCs, NKT cells and myCAFs show varying polarities across PAM50 subtypes. f) Standard deviation (SD) of the polarity (sign) of pCR Scores across all measurements. NKT cells and DCs show the highest SD.

Naturally, the results for PAM50 subtypes largely align with those based on ER subtypes, considering that ER+ samples mostly include Luminal A and Luminal B and ER-samples mostly include Basal-like and HER2-enriched. For example, myCAFs show positive pCR Scores in Luminal B and ER+ and negative pCR Scores in both Basal-like and ER-subtypes.

#### Cell types with the same polarity of pCR Scores across PAM50 sub-types

Among cancer cell types, GenMod2, GenMod4, GenMod1 and GenMod5 exhibit pCR Scores with the same polarity in at least three PAM50 subtypes. Furthermore, macrophages, CD4 T cells, RGS5 endothelials, myoepithelials and monocytes have pCR Scores with the same polarity in three PAM50 subtypes, revealing some level of consistency among breast tumor subtypes.

#### Cell types with pCR Scores only in one or two of PAM50 subtypes

Differentiated PVLs, ACKR1 endothelials, NK cells, mature luminals and GenMod6 appear in two PAM50 subtypes with nonzero pCR Scores. iCAFs and plasmablasts appear in only one PAM50 subtype.

#### Cell types exhibiting opposite polarities in pCR Scores across PAM50 subtypes

GenMod7, GenMod3, luminal progenitors, immature PVLs, myCAFs, CD8 T cells, NKT cells and DCs exhibit pCR Scores with opposite polarities between at least two PAM50 subtypes. Consistent with the previous section, DCs demonstrate positive pCR Scores in the Basal-like/HER2-enriched subtype, while displaying negative pCR Scores in the Luminal A/B subtypes. In the Basal-like subtype, myCAFs have a negative pCR Score. In Luminal B, both iCAFs and myCAFs show positive pCR Scores, aligning with findings in ER+ samples.

Despite the decreasing power due to the comparison of smaller groups, the additional stratification brings new insights as a number of associations unseen in the overall analysis appear. In addition to the aforementioned DCs, cell types such as mature luminal, GenMod3, iCAFs, CXCL12 endothelials, GenMod7 and plasmablasts appear specifically associated to response in a certain PAM50 subtype (mostly negatively)(Fig. 4). This is of particular importance in Luminal B cases, as clinically and therapeutically, they are not very distinct, sharing characteristics with both Luminal A and Basal-like subtypes. Cell types such as immature PVLs, iCAF, NKT cells and GenMod7 have negative associations for Luminal B despite positive or none in Luminal A and Basal-like PAM50 subtypes.

Thus far, we have calculated pCR Scores for seven different sample groups: the general population, two ER subtypes and four PAM50 subtypes. To assess the extent of variations in pCR Scores across them, we computed the standard deviation (SD) of their polarity across all groups, as depicted in Fig. 4 (f). Notably, GenMod2 yields an SD of zero, signifying a persistent negative pCR Score. While several cell types exhibit at least one change in their pCR Score polarity, the most pronounced variation is observed in NKT cells and DCs (also see Suppl. Fig. S13 (a)). In the subsequent section, we delve deeper into the investigation of DCs. Our focus centers on a comparison between the ER+ and ER-subtypes, given the limited number of samples, which renders a comparison among other subtypes unfeasible.

### 2.4 Further exploration of DCs

#### DC subsets have a similar abundance across ER subtypes

DCs encompass distinct subsets with both antitumor functions (conventional DCs type 1 and 2, referred to as cDC1 and cDC2) and tolerogenic functions (plasmacytoid DCs, known as pDCs) [29]. Therefore, the varying composition of DCs subsets in ER subtypes, such as a higher presence of cDC1 and cDC2 in ER- and a higher presence of pDCs in ER+ samples, can potentially account for their differing roles. To explore this possibility, we analyzed a total of 955 DCs obtained from 26 patients [8]. These cells were divided into four subsets: cDC1, cDC2, pDC and LAMP3^+^ DC. Nevertheless, the comparison of the frequency of these DC subsets shows no significant difference between ER subtypes (see Suppl. Figs S13 (b) and (c)), revealing remarkably similar composition for DCs across ER subtypes.

As the difference in abundance of subsets fails to provide an explanation, we propose that DCs play distinct roles in the TME depending on tumor subtypes. The functionality of DCs is highly context-dependent and their impact is influenced by their proximity to other cells [29–32]. For example, in Esophageal cancer, it’s not the sheer number of DCs but their proximity to epithelial cells (ECs) that impacts survival [33]. Similarly, differences in the pCR Scores of DCs may possibly be reflected in their spatial distribution, particularly with respect to ECs. To explore this possibility, we performed the following two analyses.

#### DCs are closer to ECs in ER+ samples

We investigated the differences in the spatial distribution of DCs between ER+ and ER-tumors. To accomplish this, we employed cyclic immunofluorescence (cyCIF) techniques [34], that provides an unparalleled snapshot of the proteome within the cell types in the tissue section, to generate 30-dimensional images of untreated breast tumor tissue from four patients (two samples from ER+ tumors and two samples from ER-tumors) enrolled in the NeoAva trial [35]. Utilizing Galaxy-ME for analysis [36], we identified a substantial number of individual cells in each sample. Our particular focus was on the identification of DCs, as well as ECs, to analyze their spatial distribution. Epithelial cells were classified as positive for panCK, while DCs were identified as positive for both CD45 and CD11c. All remaining cells were grouped as “other immune or stromal cells”. As Figs 5 (a) and (b) show, in the ER-sample, DCs are predominantly encircled by other immune and stromal cells, whereas in the ER+ sample, ECs also surround them. We calculated the frequency of ECs at varying distances from DCs, which is similar to Ripley’s K function [37]. Our analysis revealed that the abundance of ECs surrounding DCs was higher in the ER+ samples compared to the ER-samples, as depicted in Fig. 5 (c).

**Fig. 5:**
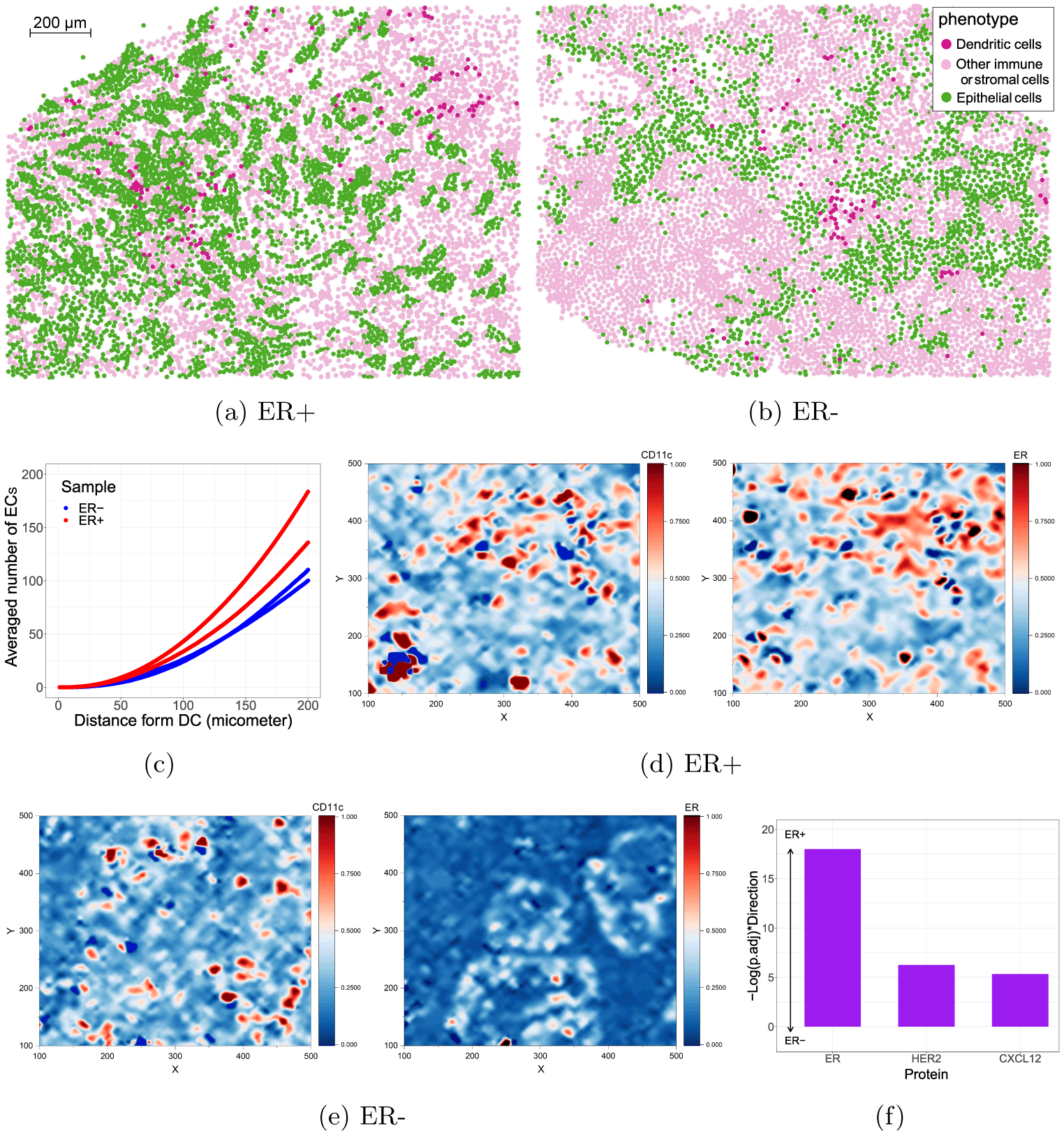
Spatial distribution of DCs in ER+ (a) and ER-(b)samples using cyCIF analysis. While DCs in the ER-sample are mostly surrounded by “other immune or stromal cells”, in the ER+ sample, they can be seen in the vicinity of ECs. c) The number of ECs in proximity to DCs was quantified, revealing that ER+ samples exhibited a higher density of ECs surrounding DCs compared to ER-samples. The normalized ER and CD11c markers density in ER+ (d) and ER-(e) samples from IMC data. f) Comparison of the correlation between CD11c (as an indicator of DCs presence) and markers ER, HER2 and CXCL12 (as indicators of ECs presence) was examined in ER+ and ER-samples, as described by [39], shows a significant difference. Both independent analyses consistently support the finding that DCs are located in closer proximity to ECs in ER+ samples compared to ER-samples.

#### Spatial correlation of markers for DCs and ECs is significantly higher in ER+ samples

According to publicly available data, to the best of our knowledge, DCs have not been identified as a distinct phenotype. However, the density of the CD11c marker has been utilized as an indicator of the presence of DCs [38]. To investigate the spatial distribution of DCs, we conducted an analysis of the spatial correlation between CD11c and three markers for ECs (ER, HER2 and CXCL12) using Imaging Mass Cytometry (IMC). Our study included 610 samples from patients enrolled in the METABRIC study [39]. Figs. 5 (d) and (e) illustrate the distribution of CD11c and ER markers in ER+ and ER-samples. It is important to note that the spatial correlation of the ER marker remains unaffected by the intensity of the marker itself, despite the significant differences observed.

As Figs. 5 (e) shows, in ER+ samples, the correlation between CD11c and ER, HER2 and CXCL12 is considerably higher, suggesting that DCs are located in closer proximity to ECs in ER+ samples compared to ER-samples. These findings provide additional support for the results presented in the previous section.

## 3 Discussion

The analysis of tumor cellular components and their impact on patient outcomes has been a topic of interest for a long time [40]. With recent advancements in scRNA-seq technologies, we now possess the capability to identify the composition of tissue samples. Using standard statistical hypothesis testing in the general population, a few cell types are significantly different between samples with and without response to NAC (see Suppl. Fig. S4). However, no cell type is significantly different when comparing pCR vs. RD for samples of specific subtypes (see Suppl. Figs S6 and S8). Only after developing an ML model and implementing our novel pipeline can we robustly identify the association between cell types and response to NAC. It is important to note that our pipeline is designed to be versatile, allowing for a robust exploration of associations between any two variables, all while comprehensively incorporating other relevant covariates.

Extending previous findings on the subtype-dependent impact of biological processes on patient outcomes in breast tumors [41], we discovered that the association of various cell (pheno)types with pCR is specific to subtypes. In the following, we discuss the main findings by cell types, starting with the ones that exhibit the most consistent results across subtypes, as shown in Fig. 4 (f):

### GenMod2

GenMod2, enriched in OxPhos and Myc, consistently exhibits negative pCR Scores in the general population as well as within all subtypes. This observation aligns with previous findings that OxPhos and Myc are potential contributors to resistance to NAC in breast cancer [42, 43].

### GenMod4

GenMod4 is enriched in G2M-Checkpoints and Mitotic-Spindle that are characteristics of proliferating cancer cells. Previous studies have shown that breast tumors with high proliferation signature scores are more likely to achieve pCR with NAC [4, 44]. GenMod4 shows the highest pCR Score in the general population and remains highly associated within all subtypes except for HER2-enriched, regardless of differences in their fractions (see Suppl. Figs S5 and S7). These findings provide additional evidence that a higher frequency of proliferating tumor cells is positively associated with pCR.

### GenMod5

Previous studies have shown that even within ER+ samples, the fraction of ER+ cells affects the chance of pCR, highlighting the role of ER+ cells in response to NAC [45]. For GenMod5, classical ER+ cancer cells, results indicate negative pCR Scores in the general population and most tumor subtypes.

### Monocytes

Monocytes serve as a bridge between innate and adaptive immune response, exerting influence on the TME through various mechanisms [46]. They display a negative pCR Score in the general population, as well as within subtypes including ER+, ER-, Luminal A, Basal-like and HER2-enriched. Monocytes consistently play a significant role in pCR across different subtypes, which represents a novel finding. This discovery should serve as an incentive for further exploration into the underlying mechanisms behind the involvement of monocytes in breast tumors.

### Macrophages

Macrophages show non-zero pCR Scores in the general population, ER+ samples, Luminal A/B and HER2-enriched subtypes. The positive pCR Score in ER+ subtype aligns with previous results [47]. In the ER-subtype, although macrophages have a significantly higher fraction, they are not associated with pCR. Furthermore, while the fraction of macrophages is significantly higher in Basal-like compared to other subtypes, it is not associated with pCR. These results suggest a potential functional difference between macrophages in Basal-like tumors and other subtypes.

### GenMod1

GenMod1 exhibits characteristics of estrogen response pathways and shows a high correlation with Fos/Jun (Activator protein 1), RTK and p53 pathways. GenMod1 has positive pCR Scores in the general population and within ER+, Luminal B, Basal-like and HER2-enriched subtypes.

### B cells

In addition to their role in antibody production, B cells can influence breast tumors through both pro- and anti-inflammatory responses [48]. However, the relationship between B cell frequency and response has not been explored. Our results demonstrate that naive B cells are negatively associated with pCR in the general population and within ER+ and ER-subtypes. Memory B cells show a negative pCR in ER-subtype.

### CD4 T cells

CD4 T cells, which can activate and regulate various aspects of innate and adaptive immunity and participate in tumor rejection [49], have been found to be positively associated with pCR in different studies [50, 51]. We also found that CD4 T cells have positive pCR Scores in the general population, as well as in ER-, Luminal A/B and Basal-like subtypes. This association may be due to CD4 T cells’ role in tumor regression during chemotherapy [52]. The frequency of CD4 T cells is significantly lower in ER+ compared to ER-subtypes within both the discovery and the validation cohorts (see Suppl. Fig. S5). Thus, the lack of association between CD4 T cells and pCR in ER+ samples may be attributed to their lower fraction. However, within Luminal A and Luminal B subtypes, CD4 T cells are positively associated with pCR, regardless of their lower fractions.

### NK cells

NK cells are innate immune cells with the ability to identify and eliminate tumor cells [53]. They were found to be positively associated with pCR in locally advanced breast tumors [54]. However, further analysis revealed that the breast TME is capable of altering the phenotype and function of NK cells [55] and their abundance is negatively associated with pCR [47]. Our results confirm the latter findings by revealing negative pCR Scores in the general population, ER+, ER- and HER2-enriched subtypes.

### PVLs

The microvascular tree consists of endothelial cells and mural cells, which include pericytes and smooth muscle cells. Mural cells not only regulate endothelial cells but can also affect tumor growth and response to chemotherapy [56, 57]. However, our knowledge of their association with chemotherapy response in breast cancer is limited [58].

Following the single-cell annotation in [8], mural cells are labeled as PVLs cells and encompass two subsets: immature PVLs and differentiated PVLs, resembling pericytes and smooth muscle cells, respectively [58]. Our results demonstrate negative pCR Scores for differentiated PVLs in the general population, both ER subtypes and Luminal A/B. Immature PVLs show negative pCR Scores in the general population, Luminal A, and Basal-like subtypes, while showing a positive pCR Score in the Luminal B subtype. Further investigation is required to understand the role of PVL subsets and how it depends on tumor subtype.

### Endothelial cells

The frequency of endothelial cells has been shown to be negatively associated with the survival of colorectal cancer patients treated with chemotherapy [59]. However, as our knowledge of endothelial phenotypes in the TME grows, the role of subsets in response to treatment remains less understood [60].

In our SM, endothelial cells are classified into four subsets, each labeled by a respective chemokine: CXCL12, ACKR1, RGS5 and LYVE1 (lymphatic endothelial). Resembling stalk-like and venular endothelial cells, ACKR1 endothelials show negative association with pCR in the general population, ER-, Luminal B and HER2-enriched subtypes. CXCL12 endothelials show positive pCR Scores in ER+, Luminal B and HER2-enriched subtypes. On the other hand, RGS5 endothelials are negatively associated with pCR in the general population, as well as in ER+, Luminal A, Luminal B and HER2-enriched subtypes.

### CAFs

CAFs play a crucial role in TME by acting as a substantial source of growth factors, cytokines and exosomes that promote tumor growth and modulate therapy responses [61–63]. In breast cancer, CAFs can promote chemoresistance by inducing stemness in breast tumor cells [64]. However, CAFs are a heterogeneous population and specific subsets involved in these functions are yet to be determined [58]. In our SM, there are two subsets of CAFs: myCAFs, enriched for myofibroblast pathways and iCAFs, which exhibit features of mesenchymal stem cells (MSCs) and inflammatory-like CAFs [65].

Our results indicate that myCAFs are negatively associated with pCR in the general population, as well as in ER- and Basal-like subtypes, with the strongest association observed in Basal-like tumors. This observation aligns with the recurring pattern of CAFs involvement in Basal-like tumors, where they, along with other stromal cells, create a reactive stroma that promotes pro-tumoral actions, including chemotherapy resistance [63]. On the other hand, in Luminal B subtype, both iCAFs and myCAFs show a positive association with pCR, indicating a subtype-specific association with response. Further analysis is needed to elucidate the mechanisms by which CAFs regulate response, but considering their diverse interactions with immune cells, they may influence NAC response by modulating the immune TME.

### Normal Epithelial Cells

Our SM includes three subsets of normal epithelial cells: myoepithelials, mature luminals and luminal progenitors. Our results reveal negative pCR Scores for myoepithelials in the general population, ER-, Luminal B, Basal-like and HER2-enriched subtypes. Although this association is new for myoepithelials, it is not unexpected given their role in breast cancer progression, where they exhibit cancer-promoting effects and contribute to tumor invasion and dissemination by producing specific cancer-promoting chemokines [66].

Mature luminals are not associated with pCR in the general population. However, in ER-positive, Luminal A and HER2-enriched subtypes, they show a positive association with pCR. The frequency of mature luminals is significantly higher in ER-positive compared to ER- and Luminal A compared to other subtypes, which might contribute to the differences in the association with pCR.

Luminal progenitors exhibit negative pCR Scores in the general population, Luminal A and HER2-enriched subtypes. However, in ER- and Basal-like subtypes, they show positive pCR Scores. The association between luminal progenitors and response to chemotherapy has not been explored so far, but tumors with bulk expression profiles resembling luminal progenitors were found to have a higher likelihood of achieving pCR [67]. Further exploration is needed to understand how luminal progenitors affect treatment response and why it differs between tumor subtypes.

### CD8 T cells

CD8 T cells, as mediators of adaptive immunity, have been extensively studied for their association with pCR and survival. While a higher fraction of CD8 T cells has been consistently associated with better survival in independent studies [68], a similar association with pCR has not been consistently reported [69]. Although the presence of CD8 T cells has been associated with a higher chance of pCR [70, 71], the analysis of CD8 T cell fraction and pCR in breast cancer has not yielded a significant association [47], warranting further investigation.

Our results reveal negative pCR Scores for CD8 T cells in the general population, ER-, and Luminal A/B, with a positive pCR Score in the HER2-enriched subtype. Such a varying behavior may explain why the association of CD8 T cells with pCR has not been consistently identified before.

### NKT cells

NKT cells, or NKT-like cells, as a subset of T cells sharing features of NK cells, work at the interface between the innate and adaptive immune systems. They can both facilitate tumor progression or exhibit antitumor activity by skewing immune responses toward tolerance or inflammation [72, 73].

Our results show a negative association between NKT cells and pCR in the general population, as well as in ER- and Basal-like subtypes. However, NKT cells also show a positive association with pCR in ER+, Luminal B and HER2-enriched subtypes. Further investigation is required to reveal the driving mechanism behind changing ploarity of pCR Score.

### DCs

DCs are antigen-presenting cells that orchestrate innate and adaptive immunity [29]. In breast tumors, DCs can contribute to immunosuppressive responses within the TME [74] or are positively associated with longer survival time [38]. Our results show a positive association between DCs and pCR in ER-, Basal-like and HER2-enriched subtypes, while a negative association is observed in ER+, Luminal A and Luminal B samples. These findings could have relevance for DC vaccines. While DC vaccines have been tested for breast samples, they have not received approval yet [75]. This might be, in part, due to their subtype-specific involvement in TME.

The composition of DC subsets, which include both antitumor functions (cDC1 and cDC2) and tolerogenic functions (pDCs), could have explained this behavior. However, our analysis of scRNA-seq data revealed a similar distribution of DC subsets across ER subtypes (see Suppl. Fig. S13). Two independent analyses, on the other hand, demonstrated a significantly different spatial distribution of DCs, with DCs being closer to ECs in ER+ samples. Considering that the proximity of DCs to ECs can influence their role in the survival of other cancer types [33], the aforementioned difference could potentially be the driving force behind the changing polarity of pCR Score of DCs in different tumor subtypes. Further investigations are required to explore how DCs are differently affected by ECs and regulate response to treatment in breast tumor subtypes.

It should be noted that we discovered that monocytes, DCs and macrophages play a central role in the response to NAC in breast tumors. These findings align with the emerging paradigm that emphasizes the importance of mononuclear phagocytes (MNPs) in the immune microenvironment, as well as in the activation and functionality of T cells [30, 76].

## 4 Methods

### 4.1 SM

The scRNA-seq data source including 100000 cells was obtained from 26 patients. Based on pseudobulk data analysis, this dataset comprises five Luminal A, three Luminal B, seven Basal-like and four HER2-enriched samples, as well as 11 ER+ and 15 ER-samples [8].

For our analysis, we randomly sampled 10,000 scRNA-seq data points. Subsequently, we employed CIBERSORTx [28] to construct an SM. The remaining 90,000 scRNA-seq cells were utilized to generate pseudobulk expression profiles for individual patients. To assess the performance of the constructed SM, we applied it to deconvolute the pseudobulk data. A comparison with the ground truth revealed a correlation of 0.70 across all cell types. Importantly, it should be noted that increasing the quantity of scRNA-seq data does not lead to enhanced performance of the SM in pseudobulk data deconvolution, as explored in the SI.

To improve the reproducibility of our results, we created ten distinct SMs. Each of these SMs was employed separately to compute cell fractions and subsequently, the estimated fractions were averaged to provide a more reliable outcome (refer to the SI for details).

### 4.2 Datasets

We divide data randomly into discovery (9 datasets, 1009 samples) and validation (6 datasets, 998 samples) cohorts ((E-MTAB-4439 [35], GSE18728 [77], GSE19697 [78], GSE20194 [79], GSE20271 [80], GSE22093 [81, 82], GSE22358 [83], GSE42822 [84], GSE22513 [85] as discovery cohort and GSE25066 [86], GSE32603 [87], GSE32646 [88], GSE37946 [89], GSE50948 [90], GSE23988 [81] as validation cohort, see SI for details). We then use our pipeline to explore the association of different cell types with pCR.

### 4.3 Machine Learning for Prediction of pCR

Through the application of deconvolution, we are able to estimate the proportions of 28 distinct cell types, while also computing the ratios of prominent cell categories, namely B cells, T cells, CAFs, PVLs, myeloids, endothelial cells, normal epithelial cells and cancer cells). Accompanying these cellular fractions are two supplementary attributes: ER status and PAM50 subtypes [16]. In our preprocessing stage, we designate positive (negative) ER statuses as 1 (0), respectively. Furthermore, we introduce dummy variables to represent the various PAM50 subtypes. This entails the transformation of the single PAM50 subtype feature into five distinct features.

We developed three prediction models: Logistic Regression, Support Vector Machines (SVM) and Random Forests, utilizing the Scikit-Learn library [91] within the Python programming language. The SVM exhibited superior performance, evaluated by a combined score of accuracy and f1 score. In the general population of the discovery cohort, the SVM outperformed the other models with an accuracy of 75.5% (±2.7%) and an f1 score of 0.429 (±0.036) in a 5-fold cross-validation. In comparison, Logistic Regression and Random Forest achieved accuracy rates of 66.0% (±2.8%) and 77.6% (±2.2%), along with f1 scores of 0.486 (±0.036) and 0.139 (±0.054), respectively (refer to SI for additional details). Consequently, the SVM was selected as the primary model for the subsequent phases of this study.

### 4.4 SHAP values

After developing a model that exhibited good predictive performance, our focus shifted toward unraveling the specific roles of different cell types in the context of pCR. In order to elucidate the contribution of individual features, we harnessed the power of SHAP (SHapley Additive exPlanations) [25], a post-hoc explanation method that calculates the marginal feature contribution to the prediction per sample.

By calculating SHAP values for each sample, we will have a set of values that describe the role of each feature in predicting pCR. Consequently, a positive SHAP value associated with a particular feature signifies its propensity to elevate the likelihood of observing a pCR outcome. Furthermore, by evaluating the absolute magnitude of the SHAP value, we gain insight into the importance of the specific feature in the prediction of pCR. This metric facilitates a ranking of features based on their respective absolute values, thereby unveiling their comparative importance in our prediction framework.

To establish a tangible measure for the influence of a cell type fraction, we conducted a linear regression analysis between SHAP values and cell type fractions. By evaluating the slope of this regression line along with its corresponding confidence interval, we can assess the overall association of the cell type fraction with pCR outcomes.

#### 4.4.1 Validation

Since SHAP values are computed based on a model, they inherently incorporate the uncertainties of the ML model [92]. To address this uncertainty, we employed a random assignment of samples into two distinct groups: the discovery cohort and the validation cohort. Subsequently, a model was trained on the discovery set, enabling the calculation of SHAP values. Subsequent to this, a linear regression was fitted to the relationship between SHAP values and their corresponding cell type fractions.

Furthermore, an independent model was trained exclusively on the validation cohort. Then the model was used to calculate SHAP values in the validation cohort. Subsequently, a linear regression line was fitted to the SHAP values, correlating them with their corresponding cell type fractions.

In instances where the slopes of the fitted lines, along with their corresponding 99.9 percent confidence intervals, exhibited the same sign for both the discovery and the validation cohorts, we considered this alignment as confirmation of a definitive, positive, or negative association between SHAP values and the fraction of the specific cell type. The same pipeline was applied to each subtype of interest.

#### 4.4.2 Calculating the pCR Score

Finally, for cell types that have successfully passed validation, the calculation of the pCR Score ensues. This entails an initial normalization of all cell type fractions to rectify for their varying abundances. Next, a linear regression model is fitted to the SHAP values plotted against their respective fractions, utilizing data from both the discovery and the validation cohorts.

Following this, the slopes of the fitted regression line are normalized, thereby assigning them the label of pCR Score. A positive (negative) pCR Score indicates a corresponding positive (negative) correlation between the fraction of a particular cell type and the likelihood of achieving pCR. Then, cell types were ordered based on their pCR Score, as illustrated in Fig. 2.

### 4.5 Spatial Omics

#### 4.5.1 cyCIF

After sample preparation and imaging (see SI for details), image analysis was performed using Galaxy-ME[36]. Individual cell quantification of each marker was performed using MCQUANT[93]. ECs and DCs phenotypes were assigned based on manually gated positivity for CKs and CD45+CD11c respectively using Scimap. The proximity between ECs and DCs was performed using the dist function of the ‘proxy’ R package.

#### 4.5.2 IMC

IMC data of [39] were downloaded from the Zenodo data repository. For samples with available ER status, the correlation between the intensity of CD11c and three ECs markers including ER, HER2 and CXCL12 was computed (see SI for details).

## 5 Data and code availability

All data and code are available at https://github.com/YounessAzimzade/XML-TME-NAC-BC (see SI for details).

## Acknowledgements

This project received funding from the European Union’s Horizon 2020 Research and Innovation Program under Grant Agreement No. 847912, BigInsight (NFR project 237718) and Integreat - The Norwegian center for knowledge-driven machine learning (NFR project 332645)

