## Supplementary Information for "Explainable Machine Learning Reveals the Role of the Breast Tumor Microenvironment in Neoadjuvant Chemotherapy Outcome"

#### 1 Deconvolution

##### 1.1 Principles of Deconvolution

Traditional bulk experiments provide an aggregate signal from thousands or even millions of cells. Deconvolution of bulk expression data refers to the computational process of estimating the proportions of different cell types that contribute to this aggregate signal. One can describe the deconvolution task as a multiple linear regression model applied to gene expression (microarray or bulk RNA seq) data, formulated as [1]:

$$\mathbf{P}_j = \mathbf{SM} \cdot \boldsymbol{\beta}_j + \boldsymbol{\epsilon} \tag{1}$$

where $\mathbf{P}_j$ is the measured expression values from tissue/tumor sample $j$, $\mathbf{SM}$ is a Signature Matrix (SM) and $\boldsymbol{\beta}_j$ is a vector of mixing fractions for sample $j$ and $\boldsymbol{\epsilon}$ represents the noise. Thus, for $P_{ij}$ being the expression of gene $i$ in sample $j$, we have: $P_{ij} = \sum_k SM_{ik}\beta_{kj}$ in which $SM_{ik}$ is the averaged expression of gene $i$ in cell type $k$ and $\beta_{kj}$ is fraction of cell type $k$ in sample $j$.

Because of the insights it offers, deconvolution has gained increasing attention and a wide range of methods have been developed [1, 2] (see [3] for a detailed list of methods and their comparisons). Here, we use CIBERSORTx, a method that is widely used [4].

##### 1.2 Creating SM

The first step in supervised deconvolution methods, such as CIBERSORTx, involves creating a suitable SM. SM is a matrix of cell-specific expression profiles and should include the cell types expected to be present in the target tissue [1, 2]. A conventional approach to creating SM is using annotated scRNA-seq data. Here, we downloaded the annotated scRNA-seq data of [5] from Gene Expression Omnibus (GEO) and Broad Institute Single Cell portal.

Following the annotation of [5], cells were categorized into nine major types. Within each of these types, the number of subsets varied based on the resolution level selected.

We chose a resolution that accurately identified all primary subsets. For instance, at this resolution, B cells were further classified into two subsets: memory B cells and naive B cells. We also made a few adjustments. Cycling PVLs, which constituted only a minor fraction (50 cells), were not included in the analysis. Additionally, cycling T cells and cycling myeloid cells were excluded as there was already a finer classification of T cells (CD4 and CD8) and myeloid (macrophages, monocytes, and DCs).

We randomly sampled 10,000 cells from all patients and used 'Create Signature Matrix' module of CIBERSORTx web tool with default settings to construct an SM (the SM and R code are available at https://github.com/YounessAzimzade/XML-TME-NAC-BC/tree/main/SignatureMatrix).

#### 1.3 Validation of SM on Pseudobulk data

To examine the performance of SM, for each patient, the remaining scRNA-seq cells was used to create a pseudobulk expression profile. The SM was then utilized to deconvolve the pseudobulk expression profile of each patient using the 'Impute Cell Fractions' module of CIBERSORTx web tool with default settings. The deconvolution results were compared with the ground truth as shown in Fig. S1 (a). The correlation between ground truth and CIBERSORTx results for all cell types was calculated. On average, the correlation between CIBERSORTx results and the ground truth was 0.70 (± 0.19) (Fig. S1 (b)), indicating a good performance.

While we included 10,000 cells, which is already more than the number required for creating a proper SM [6], we investigated whether an increase in the number of cells further affects the performance. To explore this, we selected 25,000 cells and created an SM using the Docker version of CIBERSORTx. The remaining 75000 cells were used to generate the pseudobulk data. Our results show that the performance of CIBERSORTx did not increase with the use of more scRNA-seq data (Fig. S1 (b)), as the average correlation between CIBERSORTx results and the ground truth remained at 0.67 (± 0.20).

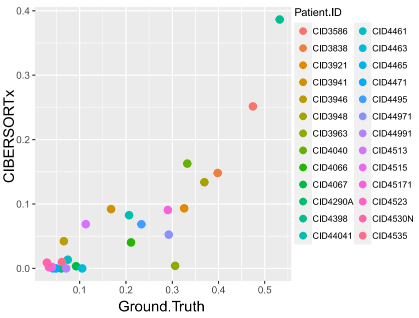


(a)

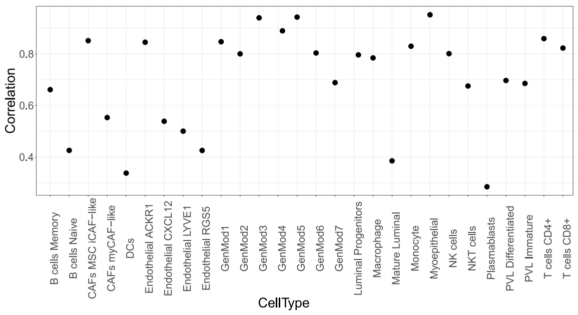


(b)

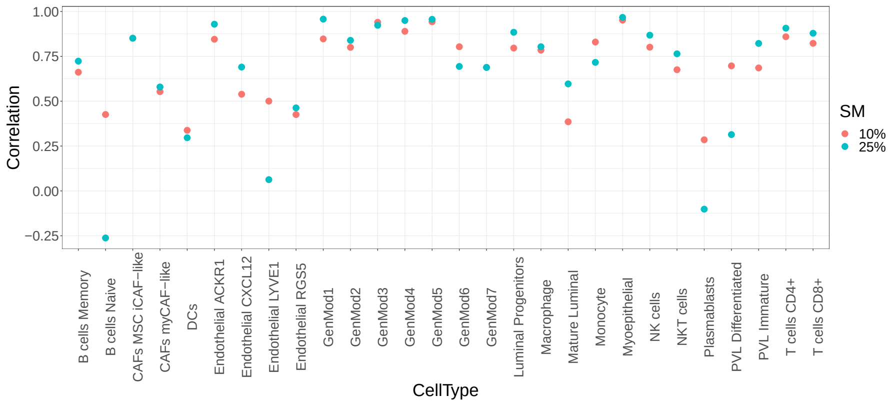


(c)

**Fig. S1**: a) Comparison of the fraction of CD4 T cells calculated by CIBERSORTx with the ground truth data for all samples shows a correlation greater than 0.8. b) We analyzed the correlation between the ground truth and CIBERSORTx results for all cell types. c) The comparison between the correlation of CIBERSORTx results and ground truth, using 10% and 25% of scRNA-seq data (10,000 and 25,000 cells, respectively), results in no significant difference in the performance.

##### 1.3.1 Uncertainty in Deconvolution

We repeat the process of creating the SM and generate a new one (SM2) that differs only in the randomness of selection of scRNA-seq data. When we use SM2 to deconvolve a dataset (in this case NeoAva [7]) and compare the calculated cell fractions with those obtained using SM1 (the initial SM), we observe a high correlation between the results. However, as depicted in Fig. S2 (a) for CD4 T cells, they are not entirely identical and carry a level of uncertainty.

This inherent uncertainty has the potential to impact the reproducibility of our findings. To address this issue, we created 10 SMs and used them in deconvolution. Subsequently, we averaged the ten fractions obtained from all ten SMs. By developing a new SM (SM11) and using it for deconvolution, the outcomes showed greater similarity, with the correlation rising to 0.94. As a result, any subsequent SM will produce cell type fractions that align more closely with the average fraction. All 10 SMs are available at https://github.com/YounessAzimzade/XML-TME-NAC-BC/tree/main/SignatureMatrix. By generating 10 SMs (SM11-20) and using them to estimate cell fractions, one can expect to observe high correlation (correlation > 0.99) when comparing these two averaged values, as shown in Fig. S2 (c).

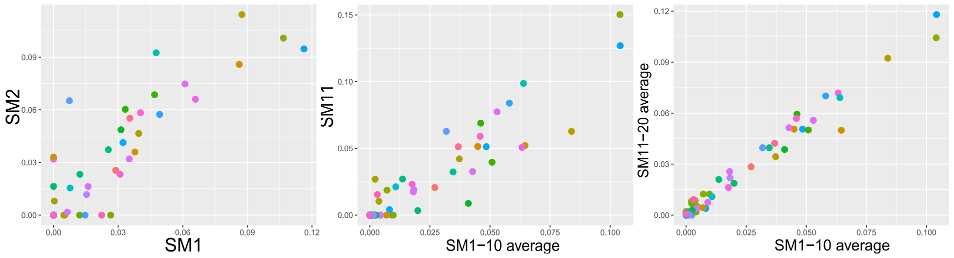


(a) (b) (c)

**Fig. S2**: a) Comparison between cell fractions estimated using two different SMs for CD4 T cells reveals that the estimated fractions are quite similar, showing a correlation of 0.91. However, it is important to note that they are not identical. b) To account for the uncertainty of our estimations, we generated 10 SMs and utilized them to estimate cell fractions. These values were then averaged to provide a more reliable estimate. Subsequently, when a new SM is introduced, and its results are compared with the averaged fractions, we observe a significant increase in similarity, with a correlation of 0.94. c) Lastly, by creating 10 SMs and comparing the averaged cell fractions, one can achieve a remarkable level of similarity, demonstrating a correlation of 0.99 between the estimations. This substantial improvement underscores the effectiveness of utilizing multiple SMs and averaging their results to enhance reproducibility.

### 2 Datasets

#### 2.1 Inclusion Criteria

Inclusion criteria was breast tumors that underwent NAC, with available bGEP microarray data and information on response to treatment (pathological complete response - pCR vs. residual disease - RD). We identified a total of 15 datasets as shown in table 1. For trials without available PAM50 classification, we used Genefu [21] R package (R code available at https://github.com/YounessAzimzade/XML-TME-NAC-BC/tree/main).

#### 2.2 Cell type fraction estimation

Each dataset was retrieved from the GEO database and subsequently, log2 normalized and scaled. In each dataset, samples with available responses to NAC (pCR

| Datasets | | | | | | |
|---|---|---|---|---|---|---|
| ID | Number of Samples | Stage | Treatment | Her2 | ER | Cohort |
| E-MTAB-4439 (NeoAva)[7] | 131 | < IV | Fluorouracil, Epirubicin and Cyclophosphamide, Taxanes (Docetaxel or Paclitaxel) + Bevacizumab | Negative | Mix | discovery |
| GSE18728[8] | 61 | II/III | Docetaxel and Capecitabine | Negative | Mix | discovery |
| GSE19697 (WASH2)[9] | 24 | | Epirubicin and Docetaxel+ Zoledronic Acid | Negative | Negative | discovery |
| GSE20194 (MDACC-MAQC)[10] | 278 | I-III | Paclitaxel, 5-fluorouracil, Cyclophosphamide and Doxorubicin | Mix | Mix | discovery |
| GSE20271[11] | 178 | I-III | Paclitaxel, 5-fluorouracil, Cyclophosphamide and Doxorubicin | Mix | Mix | discovery |
| GSE22093 (MDACC-IGR)[12] | 97 | I-III | Fluorouracil, Adriamycin, and Cytoxan | Normal | Mix | discovery |
| GSE22358[13] | 122 | I-III | Capecitabine, Docetaxel and Trastuzumab | Mix | Mix | discovery |
| GSE42822[14] | 90 | II-III | 5-fluorouracil, Epirubicin, Cyclophosphamide; Docetaxel Capecitabine | Mix | Mix | discovery |
| GSE22513 (VAN)[15] | 28 | II -III | Paclitaxel + Radiation | Mix | Negative | discovery |
| GSE25066[16] | 482 | II -III | Paclitaxel, Docetaxel with Capecitabine | Mix | Mix | validation |
| GSE32603 (I-SPY1)[17] | 138 | < IV | Taxane | Mix | Mix | validation |
| GSE32646 (Osaka)[18] | 115 | II -III | Paclitaxel, 5-fluorouracil, Epirubicin and Cyclophosphamide | Mix | Mix | validation |
| GSE37946[19] | 50 | II -III | Fluorouracil/ Epirubicin or Adriamycin/ Cyclophosphamide - Taxol | Positive | Mix | validation |
| GSE50948[20] | 156 | < IV | Doxorubicin/Paclitaxel, Cyclophosphamide Methotrexate, Fluorouracil+ Trastuzumab | Mix | Mix | validation |
| GSE23988 (USO)[12] | 61 | II -III | 5-fluorouracil, Doxorubicin, Cyclophosphamide, Docetaxel and Capecitabine | Mix | Mix | validation |

**Table 1**: bGEPs Datasets.

vs. RD) were selected. To estimate cell fractions for these samples, we utilized the 'Impute Cell Fractions' module of CIBERSORTx web tool with default settings. Cell type fractions were imputed separately using 10 SMs mentioned above and then were averaged. We obtained a total of 2007 samples from these datasets, as illustrated in Fig. S3 (a), and they are accessible on https://github.com/YounessAzimzade/XML-TME-NAC-BC/tree/main. To ensure the reliability of our findings, we compared the cell type fractions in the discovery, validation, and scRNA-seq data, revealing a good consistency in cell type fraction estimations (refer to Fig. S3 (b)).

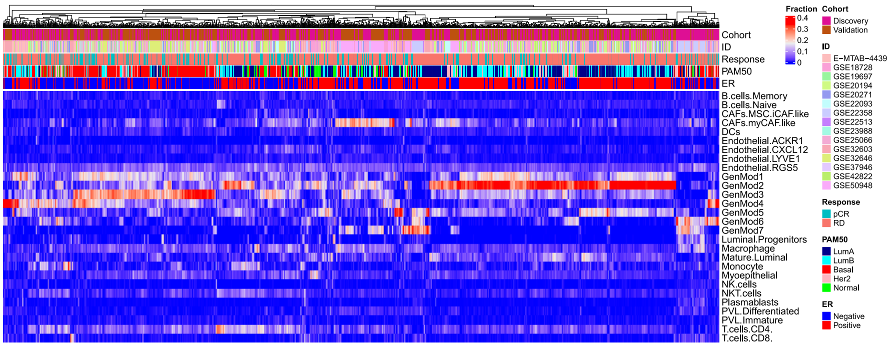


(a)

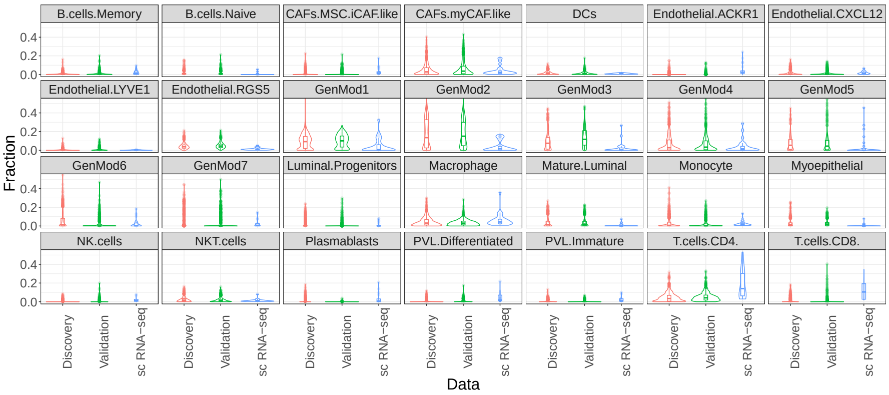


(b)

**Fig. S3**: a) Heatmap representing the estimated cell fraction, averaged over results from 10 SMs, for all samples (15 trials, 2007 samples). b) Comparison of cell fractions among discovery (1009 samples), validation (998 samples), and scRNA-seq (26 samples).

##### 2.2.1 Major cell type fractions

As previously mentioned, there exist nine major cell types. The calculation of the fractions for these major cell types involves summing up the fractions of their respective minor cell types. To illustrate, the fraction attributed to B cells would be the sum of memory and naive B cells. Plasmablasts lack a distinct minor cell type. Consequently, upon incorporating the fractions of major cell types into our ML model, a total of eight major cell types will be included.

### 3 Cell fractions comparison

#### 3.1 pCR vs. RD in general population

The comparison of cell fractions between pCR and RD in both the discovery and the validation cohorts indicates significant differences in GenMod4, mature luminal, GenMod2, and GenMod5. This observation is illustrated in Fig. S4.

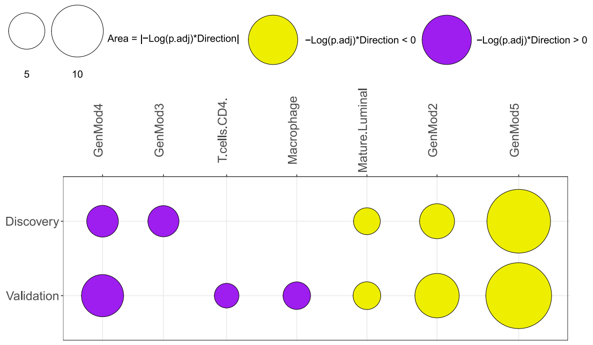


**Fig. S4**: Comparison of cell fractions in pCR vs. RD tumors in discovery and validation cohorts. Here and for the rest of the results, only cell types with significantly different fractions ($p.adj_{Bonferroni} < 0.05$) are shown.

#### 3.2 ER subtypes

##### 3.2.1 Comparisons Between ER+ and ER- samples

Comparison between cell fractions in ER+ and ER- subtypes within both discovery and validation cohorts reveals significant differences across multiple cell types. ER-subtype demonstrates higher fractions of macrophages, CD4 T cells, luminal progenitors, GenMod4, and GenMod3. On the other hand, ER+ subtypes show higher fractions of GenMod5, GenMod2, mature luminals and GenMod1.

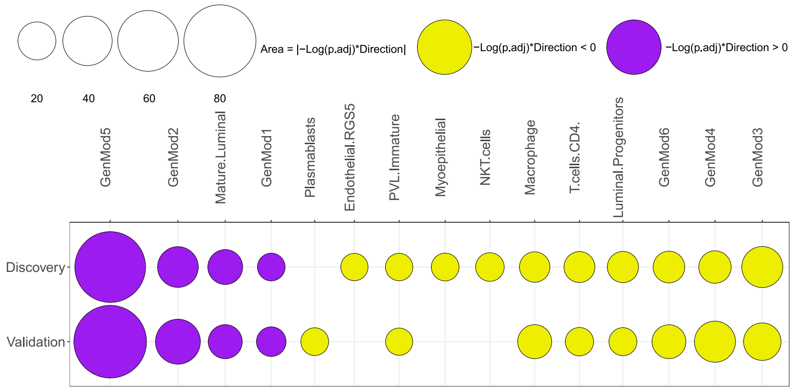


**Fig. S5**: Comparison of cell fractions in ER+ vs. ER- subtypes in discovery and validation cohorts.

##### 3.2.2 pCR vs. RD in ER subtypes

Next, for each ER subtype, we compare cell fractions in pCR vs. RD. Fig. S6 (a) shows the results for ER+ subtype and Fig. S6 (b) shows the results for ER- subtype. No cell type shows a significant difference between pCR vs. RD in both discovery and validation cohorts of either of the subtypes.

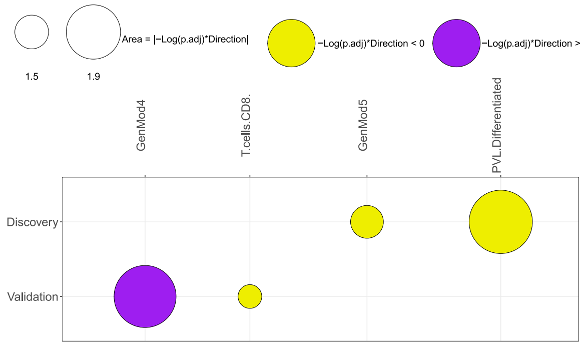


(a) ER+

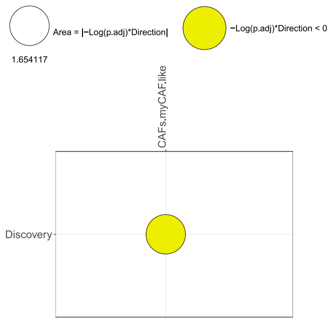


(b) ER-

**Fig. S6**: Comparison of cell fractions in pCR vs. RD tumors in discovery and validation cohorts in ER+ (a) and ER- (b) subtypes. No cell type consistently shows a significant difference in both cohorts.

##### 3.3 PAM50 subtypes

PAM50 classification includes five different subtypes: Luminal A (LumA), Luminal B (LumB), Basal-like (Basal), HER2-enriched (Her2) and Normal-like (Normal). In this work, we concentrated on the first four subtypes as the most relevant subtypes with the highest number of samples.

###### 3.3.1 Comparisons between PAM50 subtypes

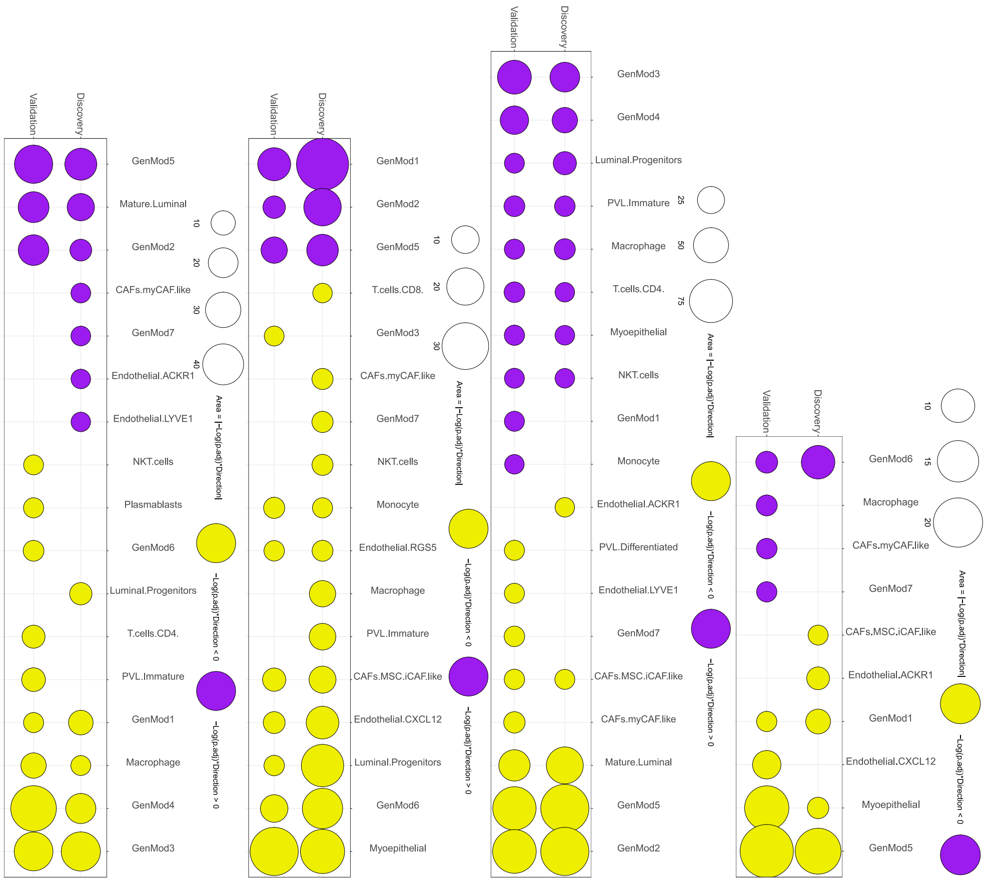


(a) LumA (b) LumB (c) Basal (d) Her2

**Fig. S7**: Comparison of cell fractions in LumA (a), LumB (b), Basal (c) and HER2 (d) vs. other subtypes in discovery and validation cohorts. The significant difference between cell fractions suggests that each PAM50 subtype has a unique microenvironment with different combinations of cancer, immune and stromal cells.

We also compare cell fractions between the PAM50 subtypes. LumA has higher fractions of GenMod5, mature luminals and GenMod2, but a lower fraction of GenMod1, macrophages, GenMod4 and GenMod3. LumB has higher fractions of GenMod1, GenMod2 and GenMod5, but lower fractions of monocytes, RGS5 endothelials, iCAFs, CXCL12 endothelials, luminal progenitors, GenMod6 and myoepithelials. Basal has higher fractions of GenMod3, GenMod4, luminal progenitors, immature PVLs, macrophages, CD4 T cells, myoepithelial and NKT cells, but lower fractions of iCAFs, mature luminals, GenMod5 and GenMod2. HER2 has higher fractions of GenMod6 and lower fractions of GenMod1, myoepithelials and GenMod5. Such substantial difference in cell fractions suggests that each PAM50 subtype has a unique microenvironment.

##### 3.3.2 pCR vs. RD in PAM50 subtypes

Comparing cell fractions between pCR vs. RD within PAM50 subtypes reveals that none of the cell types show a significant difference consistently in both cohorts.

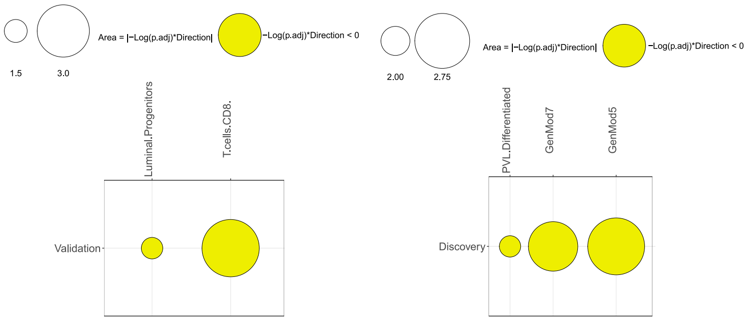


(a) LumA (b) LumB

**Fig. S8**: Comparision of cell fractions in pCR vs. RD tumors in discovery and validation cohorts in LumA (a) and LumB (b) subtypes. In Basal and HER2 subtypes no cell type showed a significant difference in pCR vs. RD.

### 4 ML Models

In the discovery cohort, we have 1009 samples with available responses to treatment (pCR or RD). Thus, we can use Supervised Machine Learning where we search for algorithms that learn training set to make predictions about future instances (test set). For such a task, there are multiple approaches, including Logistic Regression, Support Vector Machine (SVM), Random Forests, Decision Tree and K-Nearest Neighbour (KNN).

We used three different modeling approaches (Random Forests, Logistic Regression and SVM) to classify pCR vs. RD in breast tumor samples using cell fractions (28 minor + and 8 major cell types), ER status and PAM50 subtype as features or covariates (input). Models were developed using Scikit-Learn library [22] in Python. Model selection was performed based on the combination of accuracy and f1 score (accuracy in prediction of pCR multiplied by accuracy in prediction of RD) (codes for all three models are available at https://github.com/YounessAzimzade/XML-TME-NAC-BC).

For each model, hyperparameters were initially explored by GridSerachCV to find the model with the best performance. Details for the runtime of each step of the model are available in shared code. Hyperparameter tuning took 40 minutes on a Dell XPS 17 machine with Intel core i7 11th Gen processor. The SVM model provided the best performance among the three options and was used in the rest of the work.

### 5 Explainable Machine Learning

The ML methods introduce complexities and interactions that are not fully explored but only estimated. Due to these intricacies, it is often difficult to pinpoint the exact reason why an algorithm has classified an item in a certain way. As a result, many of these algorithms are referred to as a 'black-box' because their internal workings and decision-making processes remain opaque. Respectively, understanding of internal dynamics of the models becomes impossible. Explainable Machine Learning (XML) provides the means to interpret and explain how ML models arrive at their predictions or classifications. By deciphering the inner workings of ML models, XML enables the identification of critical features driving model predictions, aiding in knowledge discovery.

#### 5.1 SHAP values

Among XML approaches, SHAP (SHapley Additive exPlanations) [23] is widely used. SHAP is a method to measure the importance of features in making a classification of a sample. To do so, SHAP relies on Shapley's values. Named in honor of Lloyd Shapley, who introduced it in 1951 and won the Nobel Memorial Prize in Economic Sciences for it in 2012, the Shapley value is a method to distribute the total gains to the players, assuming that they all collaborate. The SHAP method draws on cooperative game theory, leveraging the concept of Shapley values to quantify the contribution of each feature to the prediction outcome. According to the Shapley value, the amount that features $i$ is given in a prediction of a ML model is calculated as:

$$\phi_i = \sum_{S \subseteq F \setminus \{i\}} \frac{|S|!(|F| - |S| - 1)!}{|F|!} (f_{S \cup \{i\}}(x_{S \cup \{i\}}) - f_S(x_S)) \tag{2}$$

where $f_{S \cup \{i\}}$ is trained model with that feature present, and $f_S$ is another model trained with the feature withheld. Then, predictions from the two models are compared on the current input $f_{S \cup \{i\}} - f_S$, where $x_S$ represents the values of the input features in the set $S$. Since the effect of withholding a feature depends on other features in the model, the preceding differences are computed for all possible subsets that lack feature $i$, $S \subseteq F \setminus \{i\}$ [23].

By computing the Shapley values for all features, SHAP assigns a fair share of importance to each feature based on its impact on predictions. This approach helps to unravel the black-box nature of ML models, providing insights into which features are driving the model's decisions. With SHAP, it becomes possible to identify influential factors.

SHAP provides different model-specific and model-agnostic explainers. Within SHAP Python package, we use KernelExplainer which is an implementation of Kernel SHAP and a model agnostic method to estimate SHAP values for any model. Kernel SHAP uses a special weighted linear regression to compute the importance of each feature. The computed importance values are Shapley values from game theory and also coefficients from a local linear regression.

To perform a SHAP analysis, one needs a trained model, data on which the SHAP values will be calculated and the background data. Background data serves as a reference distribution against which the model's behavior is compared. By using a background dataset, SHAP establishes a baseline against which the contributions of different features are measured. The background data represents the "average" or "typical" input instance, which allows SHAP to quantify how each feature deviates from this baseline. This comparison helps in understanding the importance of features in driving predictions [23].

To determine the impact of a feature, that feature is set to "missing" and the change in the model output is observed. Since most models aren't designed to handle arbitrary missing data at test time, SHAP simulates "missing" by replacing the feature with the values it takes in the background dataset. So if the background dataset is a simple sample of all zeros, then SHAP would approximate a feature being missing by setting it to zero [23]. One dataset can simultaneously be used as the background data and the data for which SHAP values are being calculated and throughout this work, we follow this approach. Python scripts for ML and SHAP are available at https://github.com/YounessAzimzade/XML-TME-NAC-BC/tree/main/pCRScores/GeneralPopulation.

#### 5.2 Uncertainty in SHAP values

SHAP values can be interpreted as a transformation in representation, wherein disparities in cell fractions are effectively delineated and distinguished through the employment of the ML model. However, it is important to note that this transformation remains susceptible to uncertainties inherent in the underlying ML model [24].

A manifestation of this uncertainty arises when the data is partitioned into three segments, and the model yielding the optimal performance on two-thirds of the data (referred to as Model1, Model2, and Model3) is selected. Subsequently, this chosen model is utilized for the computation of SHAP values. Upon comparing these SHAP values, a discernible variation comes to light, underscoring a degree of dissimilarity. For a visual depiction, refer to Figure S9, which presents a graphical representation of SHAP values in relation to the fraction of GenMod5, employing three distinct models.

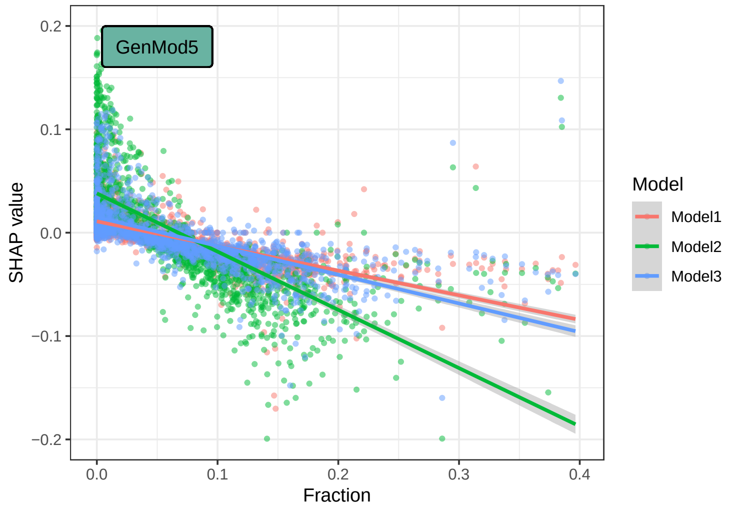


**Fig. S9**: SHAP values vs. fraction of GenMod5 using three models. Each point represents a sample. Identical samples show different SHAP values because of differences in ML models.

To mitigate these uncertainties, we divide the data into two distinct groups, one for discovery and one for validation. This enhances the robustness of the SHAP values by accounting for potential variations arising from the uncertain nature of the ML model.

#### 5.3 Validation

We fine-tune the hyperparameters to discover the model with the best performance. Subsequently, we calculate the SHAP values utilizing the best-performing model (calculated SHAP values for general populations of both cohorts are available at https://github.com/YounessAzimzade/XML-TME-NAC-BC/tree/main/pCRScores/GeneralPopulation).

To gain an overview of which features hold the most significance for the model's prediction, we sort features based on average absolute SHAP values over all samples, as shown in Fig. S10(a). Given our focus on TME, we explore the association between cell fractions and SHAP values. We plotted SHAP values against normalized cell fractions, as illustrated in Fig. S10(b). This reveals that not all cell types exhibit a clear association between SHAP values and cell fractions. Specifically, for some cell types, their 0.999 confidence intervals include zero. Consequently, these particular cell types do not progress beyond this stage of analysis

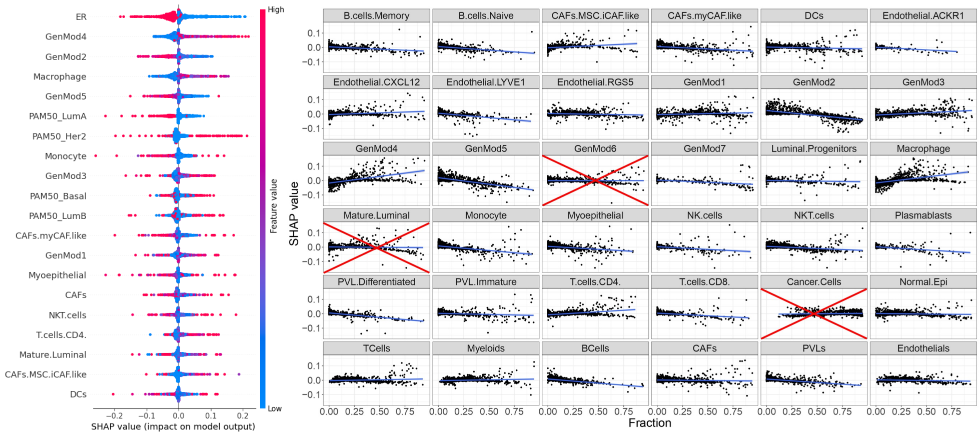


(a) (b)

**Fig. S10**: a) The top 20 features ranked based on the average absolute SHAP values of the discovery cohort. The color in the plot represents the feature value, with red indicating high values and blue indicating low values. As expected, features such as ER status and PAM50 subtypes alongside different cell fractions appear among the most important features. We are interested in cell types, thus we only look into their SHAP values. Additionally, we are interested if there is a relation between cell type fraction and their SHAP value. Thus we fit a linear line and if the 0.999 confidence interval doesn't include zero, one can argue that there is an association between SHAP values and cell fractions.

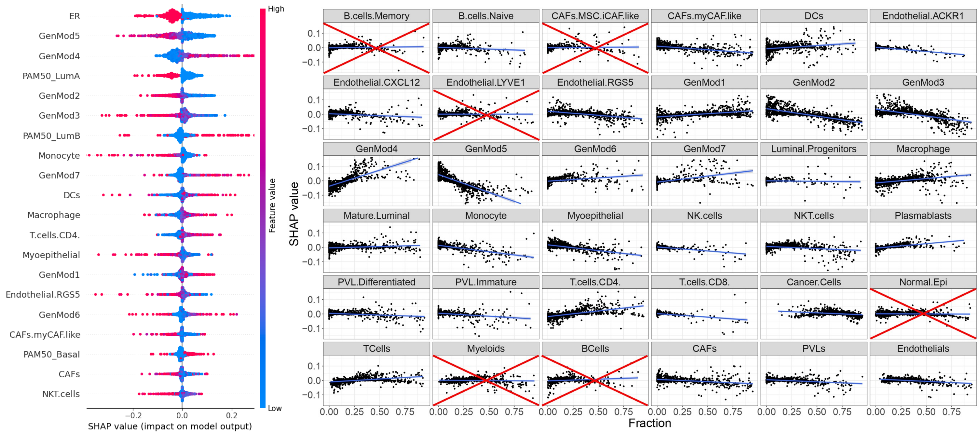


(a) (b)

**Fig. S11**: a) Top 20 features ranked based on the average absolute SHAP values of the validation cohort. b) Cell types that don't show an association are removed.

We then use a similar pipeline and calculate SHAP values in the validation cohort. By comparing Figs S10 (a) and S11 (a), one can see that the results are relatively similar. ER status, for example, is the most important feature in the prediction of pCR in both cohorts. However, to have a concrete comparison, we take all cell types that show a clear association between SHAP values and pCR.

#### 5.4 Calculating pCR Score

When we analyze the SHAP values of all cell types, some of these have already failed our initial validation. Among the cell types that successfully pass this preliminary assessment, those lacking a consistent association with pCR in both cohorts will be further excluded (see Fig. S12 (a)). For the remaining cell types, we construct a regression line to model the relationship between SHAP values and cell fraction.

To facilitate meaningful comparisons and minimize the impact of extreme outliers, we applied a two-step normalization process. Percentile-Based Scaling: Initially, slopes were scaled based on the 95th percentile. This operation ensured that the value corresponding to the 95th percentile of our data was normalized to 1, while all other values were scaled proportionally. Value Clipping: Post-scaling, we observed that some scaled values, due to their original magnitude, exceeded the desired range of -1 to 1. To address this, we applied a constraint to our scaled data. Values exceeding 1 were set to 1, and those below -1 were set to -1. As a result of this combined approach, our normalized data was effectively bound within the range of -1 to 1.

Subsequently, we arrange the cell types in descending order according to their respective pCR Scores, as shown in Fig. S12 (b). The R code for implementing this pipeline is available at https://github.com/YounessAzimzade/XML-TME-NAC-BC.

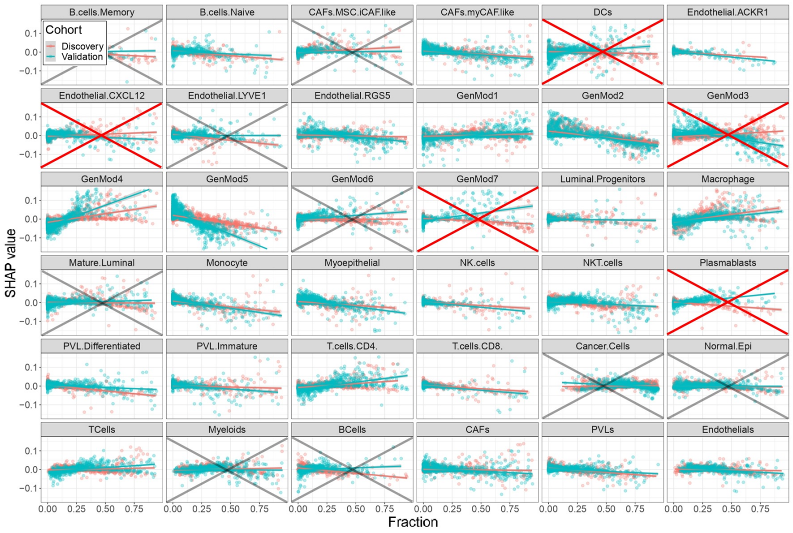


(a)

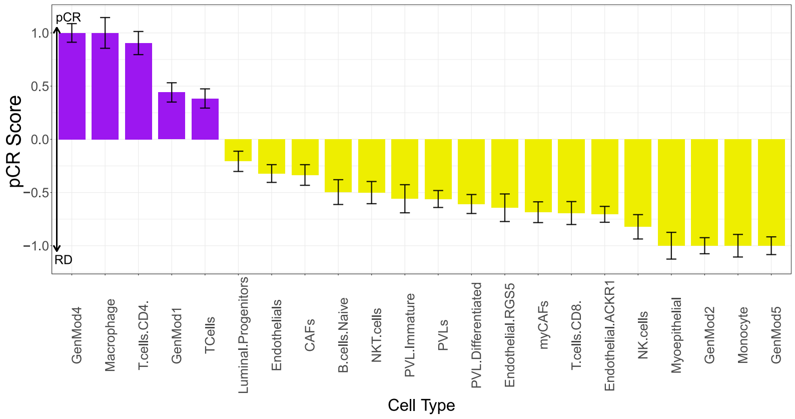


(b)

**Fig. S12**: a) Cell types that lack a discernible association between SHAP values and fractions in either of the cohorts have already been excluded. Furthermore, any cell types that fail to exhibit a consistent association across both cohorts are removed at this stage. b) Among the cell types that remain, we perform linear regression on samples from both cohorts. We then normalize coefficients as pCR Score and sort cell types based on their pCR Scores.

##### 5.4.1 pCR Score of major cell types

We have encompassed eight major cell types in our study. However, since they largely adhere to their prevailing subset, we have excluded them from the final analysis. For instance, among T cells, CD4 T cells exhibit a more pronounced correlation with pCR, and the pCR Score of T cells corresponds to this correlation. as shown in Fig. S12 (b).

### 6 Further exploration of DCs

To evaluate the variability in pCR Score from the different calculations we've performed, we also determined its standard deviation (SD) as depicted in Fig. S13 (a). DCs show the highest value of SD.

#### 6.1 DCs subsets

Following available annotations [5], cells were divided into four subsets: cDC1, cDC2, pDC, and LAMP3$^+$ DC. We compare the frequency of subsets of DCs in ER+ vs. ER- tumors. The comparison reveals that ER- samples have higher frequencies of all DCs subsets (no significant difference). Upon accounting for discrepancies in the overall number of DCs, the relative proportions of subsets across ER subtypes exhibited greater similarity. This observation suggests a more uniform distribution of DC subset frequencies across ER subtypes (data Available at https://github.com/YounessAzimzade/XML-TME-NAC-BC/tree/main/DCs/scRNAseq)

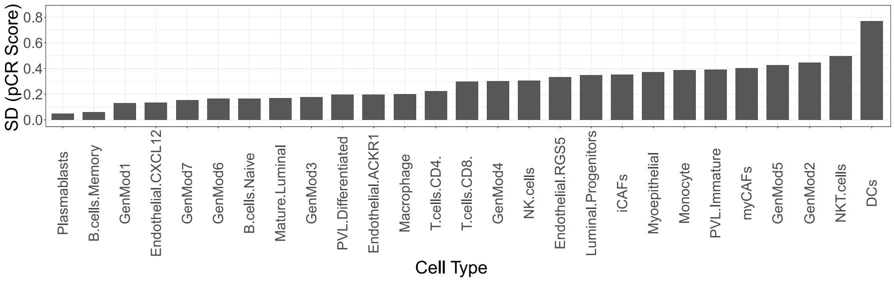


(a)

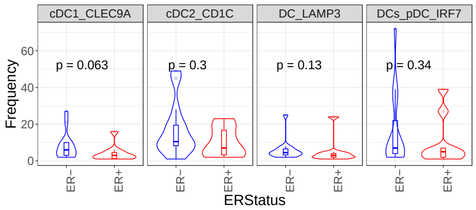


(b)

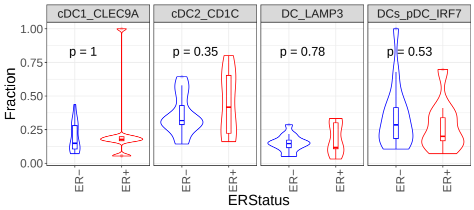


(c)

**Fig. S13**: a) Standard deviation of the pCR Score across all subtypes. Similar to the SD of polarity, DCs show the highest variation across subtypes. b) Frequencies of DC subsets in ER subtypes do not exhibit any significant differences. c) Upon normalization for overall frequency, the average fractions of DC subsets become remarkably similar across ER subtypes.

#### 6.2 cyCIF

Cyclic immunofluorescence (cyCIF) is a cutting-edge technique to analyze multiple cellular proteins simultaneously with high spatial resolution. Using several sequential rounds of immunofluorescence staining and quenching, it allows for detection of more than 40 proteins on a single FFPE slide.

In this study, the samples were processed as follows: FFPE slides were baked overnight at 55°C followed by 30 minutes at 65°C, before deparaffination. Antigen retrieval was performed using a Cuisinart pressure cooker (model CPC-600). Briefly, slides were emerged in a coplin jar filled with pH6 Citrate buffer. The pressure cooker was set on high for 4 minutes, and slides were incubated in the pressure cooker for 20 minutes before releasing the pressure. Slides were then rinsed once into warm deionized water and incubated for 15 minutes in warm pH9 Tris/EDTA antigen retrieval buffer. After blocking the slides for 30 minutes at room temperature in a solution of PBS containing 10% NGS (Vector) and 1% BSA (Sigma), coverslips were mounted using slowfade Gold DAPI (Life Technologies) mounting media and slides were scanned at 10X magnification using the Axioscan Z1 fluorescence slide scanner (Zeiss). Autofluorescence of each tissue was acquired.

After imaging, the coverslips were removed by soaking the slides in a vertical position within a staining dish filled with PBS. Slides were then incubated with the first round of four antibodies, directly labelled with different Alexa-Fluor (AF) molecules (AF488, AF555, AF647 and AF750) for 2 hours at room temperature. After acquiring an image of the staining, the fluorescence molecules were quenched using a solution of 3% peroxide and 20 mM NaOH in PBS. Subsequent rounds of staining and quenching were then performed on each slide until all antibodies were probed. In this study antibodies against CD45, CD11c and pan-cytokeratin (CKs) is reported as shown in Fig. S14.

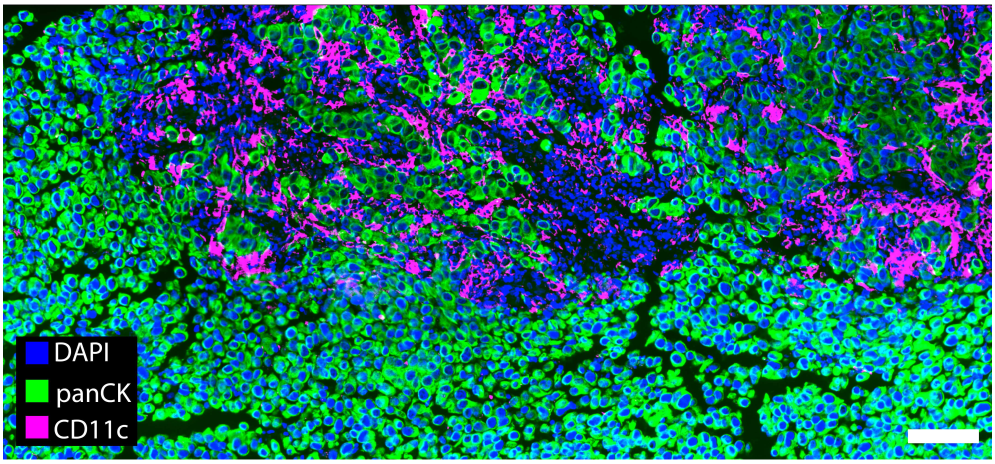


(a) ER+

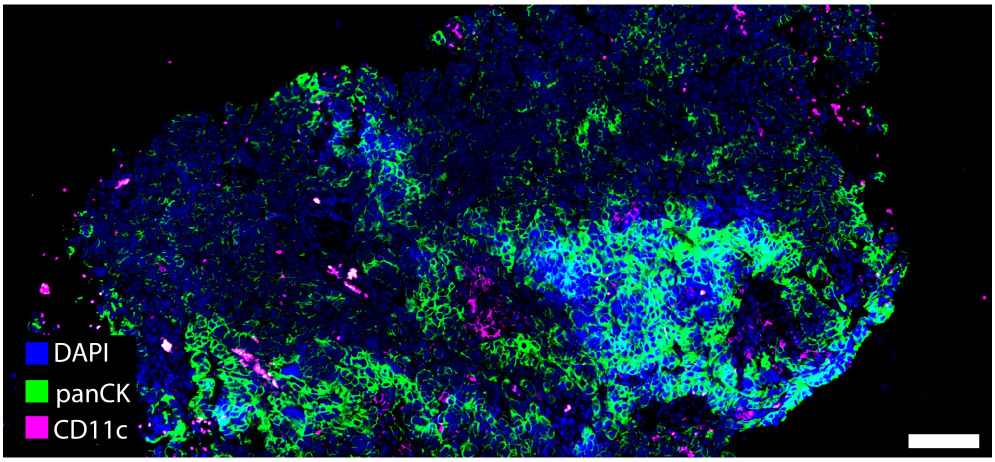


(b) ER-

**Fig. S14**: cyCIF slide for ER+ (a) and ER- (b) samples (scalebar is 100 micrometer).

Image analysis was performed using Galaxy-ME [25]. In short each image acquired during cyCIF was corrected for illumination using ImageJ BaSiC[26], then registered based on DAPI staining using ASHLAR[27] followed by whole-cell segmentation based on DAPI and cell membrane markers (ECadherin, CD45 and CKs) using Mesmer[28]. Individual cell quantification of each marker was performed using MCQUANT[29]) for which the signal intensities were adjusted for autofluorescence background (SBR). Epithelial (EC) and dendritic cell (DC) phenotypes were assigned based on manual gated positivity for CKs and CD45+CD11c respectively using Scimap and visualized using Vitessce[30].

The calculation of proximity between ECs and DCs was carried out using the exported AnnData from Galaxy-ME. To achieve this, the dist function from the 'proxy' R package was employed. Subsequently, the number of ECs within specific distances from DCs was computed. This measurement is similar to Ripley's K function [31], and was applied across two distinct populations (ECs and DCs). The relevant code and data can be accessed at https://github.com/YounessAzimzade/XML-TME-NAC-BC/tree/main/DCs/SpatialOmics/cycIF.

#### 6.3 IMC

We downloaded the imaging mass cytometry (IMC) data of [32] from the Zenodo data repository. Samples with available ER status were selected and the correlation between the intensity of CD11c and three ECs markers including ER, HER2 and CXCL12 was computed in each sample. As a control, we also calculated the spatial correlation of the CD16, the top marker for myeloids, with the aforementioned ECs markers. A comparison between these correlations in ER+ vs. ER- shows that CD11c correlations with all three markers are higher in ER+, suggesting that DCs are closer to ECs in ER+ samples. On the other hand, the correlation of CD16 with ER shows no difference between ER+ vs. ER- samples. Additionally, CD16 correlations with HER2 and CXCL12 markers show a smaller difference in ER+ vs. ER- samples. Code and data available at https://github.com/YounessAzimzade/XML-TME-NAC-BC/tree/main/DCs/SpatialOmics/IMC.

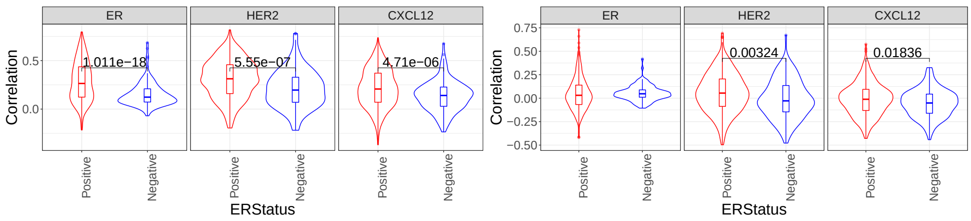


(a) CD11c (b) CD16

**Fig. S15**: a) Spatial correlation of CD11c and three EC markers. There is a significant difference between the correlations across ER subtypes. b) Spatial correlation of CD16, first marker for myeloids [32], and EC makers. The spatial correlation of DC16 with ER maker shows no difference between ER subtypes.
